# Glial insulin regulates cooperative or antagonistic Golden goal/Flamingo interactions during photoreceptor axon navigation

**DOI:** 10.1101/2020.01.23.916403

**Authors:** Hiroki Takechi, Satoko Hakeda-Suzuki, Yohei Nitta, Yuichi Ishiwata, Makoto Sato, Atsushi Sugie, Takashi Suzuki

## Abstract

Transmembrane protein Golden goal (Gogo) interacts with the atypical cadherin Flamingo to direct R8 photoreceptor axons in the *Drosophila* visual system. However, the precise mechanisms underlying Gogo regulation during columnar- and layer-specific R8 axon targeting are unknown. Our studies demonstrated that the insulin secreted from surface and cortex glia switches the phosphorylation status of Gogo, thereby regulating its two distinct functions in this process. Nonphosphorylated Gogo mediates the initial recognition of the glial protrusion in the center of the medulla column, whereas phosphorylated Gogo suppresses horizontal filopodia extension by counteracting Flamingo to maintain one axon to one column ratio. Later, Gogo expression ceases during the midpupal developmental stage, thus allowing R8 filopodia to extend vertically into the M3 layer. These results demonstrate that the long- and short-range signaling between the glia and R8 axon growth cones regulates growth cone dynamics in a stepwise manner, and thus shape the entire organization of the visual system’s functional neuronal circuit.

## Introduction

During development, well-defined synaptic connections are formed in the brain between specific neurons to facilitate higher-order information processing. Synapses are often arranged into structures that reflect the functional organization of synaptic contacts (Huberman et al., 2010; Luo and Flanagan, 2007; Sanes and Yamagata, 2009). Each brain layer receives discrete axonal inputs that carry specific information. Therefore, external inputs dissolve into distinct modules in the brain. In the visual system, photoreceptors connect to columns located around the target region, thereby preserving the spatial relationships between the visual world and its representation in the brain (Huberman et al., 2010; Sanes and Zipursky, 2010). Layers separate the brain into horizontal planes, whereas columnar units group the axons into bundles that are perpendicular to the layers (Clandinin and Zipursky, 2002; Mountcastle, 1997; Sanes and Zipursky, 2010). The integration of the individual column and layer processes enables the modular processing of perceived information. Thus, specific layer–column axonal targeting to unique synaptic partners is a fundamental step in the complex formation of functional neuronal networks inside the brain (Huberman et al., 2010; Luo and Flanagan, 2007; Millard and Pecot, 2018; Neriec and Desplan, 2016).

The *Drosophila* visual system is an attractive model for studying the formation of the functional organization of synaptic connections because its optic ganglion has a layered and columnar structure (Hadjieconomou et al., 2011; Millard and Pecot, 2018; Sanes and Zipursky, 2010). The visual system of the adult *Drosophila* consists of the compound eye and four optic ganglia (in order: lamina, medulla, lobula, and lobula plate). The compound eye is composed of an array of approximately 800 ommatidia, each containing 8 photoreceptor cells (R cells, R1–R8) arranged in a stereotypic pattern. R7 and R8 axons project to the second optic ganglion, namely, the medulla. The medulla is subdivided into columnar units and 10 distinct layers. R7, R8, and Mi1 axons elongate into the medulla at the earliest stage. They function as the pioneering axons during the formation of the medulla columns, which are comprised of approximately 100 different axons (Trush et al., 2019). R8 extends its axon to a single medulla column, followed by a single R7 axon. Eventually, R8 targets the M3 layer of the medulla, whereas R7 targets the M6 layer. Across development, the R8 neurons undergo three stages of axonal targeting (Akin and Zipursky, 2016; Hadjieconomou et al., 2011). First, single R8 axons project to a single column and form a horseshoe-shaped terminal that encircles the medulla columnar center (step 1). Second, the R8 axons remain at the medulla neuropil surface without bundling with each other (step 2). Third, R8 axons extend filopodia to target the M3 layer (step 3). Many studies have detailed the molecular mechanisms that underlie the layer-specific targeting of R neurons (Akin and Zipursky, 2016; Hadjieconomou et al., 2011; Hakeda-Suzuki and Suzuki, 2014; Hakeda-Suzuki et al., 2017; Kulkarni et al., 2016; Mencarelli and Pichaud, 2015; Millard and Pecot, 2018; Ozel et al., 2015). However, little is known about the formation of the medulla columnar structure.

Previous work in our lab identified a single transmembrane protein, Golden goal (Gogo), by a large-scale screen to search for genes that control R axon pathfinding (Berger et al., 2008). Functional studies have revealed that Gogo, with the atypical cadherin Flamingo (Fmi), guides R8 axons to the M3 layer (Hakeda-Suzuki et al., 2011; Senti et al., 2003; Tomasi et al., 2008). Gogo and Fmi colocalization is essential for this function. The R8 axons of *gogo* or *fmi* single mutants exhibit similar phenotypes, including defects in the axonal array due to the irregular distances between axons and the difficulty in targeting the M3 layer. Furthermore, the dephosphorylated state of a triplet Tyr-Tyr-Asp (YYD) motif in the Gogo cytoplasmic domain is important to R8 axon targeting (Mann et al., 2012). However, when the YYD motif is phosphorylated, Gogo appears to interfere with the ability of the R8 axon to target the M3 layer. The *Drosophila* insulin receptor (DInR), a tyrosine kinase receptor, is one of the kinases that phosphorylate the YYD motif of Gogo (Mann et al., 2012). A growing number of recent studies have revealed the functional involvement of DInR in nervous system development (Fernandes et al., 2017; Rossi and Fernandes, 2018; Song et al., 2003). Therefore, DInR may be one mechanism through which Gogo and Fmi regulate R8 axon pathfinding. Because Gogo and Fmi are conserved across C. elegans to humans, elucidating their role in development in Drosophila can greatly enhance our understanding of molecular mechanisms of development in higher-order species.

The current study was able to examine stepwise R8 axonal targeting events across development by following protein localization and by specifically controlling Gogo and Fmi levels in R8 axons. In step 1, Gogo and Fmi cooperated in guiding the R8 growth cone to its correct place inside the column. In step 2, Gogo was phosphorylated by the glial insulin signal and began to counteract Fmi to repress filopodia extension. In step 3, R8 axons only expressed Fmi, which directed them to the M3 layer. These results indicate that the glial insulin signal controls Gogo phosphorylation, thereby regulating growth cone dynamics, including the formation of the horseshoe shape and filopodia extension. Overall, this regulates axon–column and axon–axon interactions. Gogo possesses an interesting property wherein the phosphorylation states maintain two separate axon pathfinding functions. This is an economical strategy for increasing protein functions when there are a limited number of genes. As a result, this mechanism maintains the regular distance between R8 axons and enables the ordered R8 axonal targeting of the column.

## Materials and Methods

### Fly strains and genetics

Flies were kept in standard Drosophila media at 25°C unless otherwise indicated. The following fly stocks and mutant alleles were used: sens-FLP, 20C11FLP, GMR-(FRT.Stop)-Gal4 (Chen et al., 2014); gogo[H1675], gogo[D1600], UAS-GogoT1, ato-Δmyc, GMR-GogoΔN-D, GMR-GogoΔN-E, GMR-GogoΔN-G, GMR-GogoΔN-H, UAS-GogoFL-myc, UAS-gogoΔC-myc, UAS-gogoΔN-myc (Tomasi et al., 2008); UAS-GogoΔC, <gogo<, <fmiN<, fmi[E59], UAS-Fmi, UAS-Fmi ΔC (Hakeda-Suzuki et al., 2011); UAS-GogoFL-P40, UAS-GogoFFD-P40, UAS-GogoDDD-P40, GMR-GogoFFD-myc, GMR-gogoDDD-myc (Mann et al., 2012); UAS-add1-myc, hts[null], (Ohler et al., 2011); sens-lexA, LexAop-myrTomato, bshM-Gal4, UAS-myrGFP (Trush et al., 2019); GMR-Rho1 (Hariharan et al. 1995); dilp7-Gal4 (Yang et al., 2008); dlip4-Gal4 is a gift from Dr. Pierre-Yves Plaçais (CNRS France).

Following stocks used in this study are available in stock centers: UAS-FRT-stop-FRT-mcd8GFP, loco-Gal4, Act-Gal4, sensGal4, R85G01Gal4, R25A01Gal4, Mz97Gal4, UAS-stinger, Rh6-mCD8-4xGFP-3xmyc, Rh4-mCD8-4xGFP-3xmyc, gogo-Gal4, OK107-Gal4, UAS-dicer2, UAS-40D, tub-gal80[ts], UAS-FLP, UAS-mcd8GFP, UAS-myrRFP, UAS-nlsGFP, UAS-shi[ts1], UAS-htsRNAi,, UAS-hob RNAi, UAS-dlip1 RNAi, UAS-dlip2 RNAi, UAS-dlip3 RNAi, UAS-dlip4 RNAi, UAS-dlip5 RNAi, UAS-dlip6 RNAi, UAS-dlip7 RNAi, UAS-dlip8 RNAi, dlip1-Gal4, dlip2-Gal4, dlip3-Gal4, dlip5-Gal4, UAS-Fz RNAi, UAS-Fz2 RNAi, UAS-dsh RNAi, UAS-Gq RNAi, UAS-Go RNAi, UAS-GsRNAi, UAS-Gi RNAi, UAS-Gf RNAi, UAS-cta RNAi (BDSC); dilp6-Gal4 (DGRC); UAS-gogoRNAi UAS-fmiRNAi (VDRC). The following fly strains were generated in this work: gogo-FSF-GFP, fmi-FSF-mcherry, gogoΔGOGO1, gogoΔGOGO2, gogoΔGOGO3, gogoΔGOGO4, gogoΔCUB, gogoΔTSP1, gogoFlpstop. The specific genotypes utilized in this study are listed in Table S1.

### Generation of Gogo-FsF-GFP and Fmi-FsF-GFP knock-in allele

Gogo-FsF-GFP and Fmi-FsF-GFP knock-in allele was generated by CRISPR/Cas9 technology (Kondo and Ueda, 2013). A knock-in vector containing the homology arms, the flip-out cassette with GFP (FRT-stop-FRT-GFP), and the red fluorescent transformation marker gene (3xP3RFP) was generated as described previously (Trush et al., 2019). The oligo DNAs used for amplification of Gogo and Fmi fragments and creating gDNA are listed in Table 2. A gRNA vectors were injected to eggs of yw; attP40{nos-Cas9}/CyO or y1 w1118; attP2{nos-Cas9}/TM6C, Sb Tb together with the knock-in vector. The precise integration of the knock-in vector was verified by PCR and sequencing.

### Generation of gogo mutants deleting a specific domain

gogoΔGOGO1, gogoΔGOGO2, gogoΔGOGO3, gogoΔGOGO4, gogoΔCUB, gogoΔTSP1 mutants were generated by CRISPR/Cas9 technology (Kondo and Ueda, 2013). A part of gogo gene deleting a specific domain were amplified by overlapping PCR. Single or multiple gDNA vectors were created and cloned into pBFv -U6.2 vector. The DNA oligos used for cloning and creating the gDNA are listed in Table 2.

### Generation of Gogo FRTStop mutant

GogoFlpStop mutant was generated by replacing gogo intronic MiMIC cassette (BDSC; 61010) with the FlpStop cassette using C31 integrase (Hu et al., 2011). The FlpStop cassette is a gift from Dr. Thomas R Clandinin.

### Immunohistochemistry and imaging

The experimental procedures for brain dissection, fixation and immunostaining as well as agarose section were as described previously (Hakeda-Suzuki et al., 2011). The following primary antibodies were used: mAb24B10 (1:50, DSHB), rat antibody to CadN (Ex#8, 1:50, DSHB), mouse antibody to Repo (8D12, 1:20, DSHB) mouse antibody to myc (4E10, 1:100, Santa Cruz), rabbit antibody to RFP (1:500 ROCKLAND), rabbit antibody to GFP conjugated with Alexa488 (1:200, Life technologies). The secondary antibodies were Alexa488, Alexa568 or Alexa633-conjugated (1:400, Life technologies). Images were obtained with Nikon C2^+^ and A1 confocal microscopes and processed with Adobe Photoshop and Illustrator.

Live imaging was done according to (Özel et al., 2015). Images were obtained with Zeiss LSM880NLO + COHERENT Chameleon Vision.

## Results

### Gogo expression, but not Fmi expression, ceases around the midpupal stage

During development, Gogo and Fmi proteins are expressed broadly and dynamically in photoreceptors and the optic lobe. To monitor the precise expression and localization patterns of Gogo and Fmi proteins during R8 axonal targeting, knock-in flies that tag the desired proteins in a cell-specific manner with GFP or mCherry were generated using the CRISPR/Cas9 system (Chen et al., 2014; Kondo and Ueda, 2013; Sander and Joung, 2014). The use of these flies allowed the observation of endogenous R8 axon-specific Gogo and Fmi localization across the developmental stages between the third instar larvae and adulthood (Figure 1). Gogo protein was strongly expressed in the tip of R8 axons during developmental steps 1 and 2 (Figure 1C–1E). Contrary to previous hypotheses (Hakeda-Suzuki et al., 2011), Gogo protein was not present during step 3, when R8 axons filopodia elongate toward the deeper medulla layers (Figures 1F, 1G and S1). Conversely, Fmi–mCherry expression in R8 axons was observed throughout the development stages (Figures 1H–1K). Fmi was localized in the R8 axon tip, including thin filopodia structures during step 3, when Gogo expression was not present (Figure 1K). Gogo and Fmi protein localization in the R8 axon tip during step 1 had several characteristics (Figures 1M–1O). Gogo–GFP signal was rather weak in the filopodia but accumulated at the rim of the horseshoe-shaped axon terminal that encircled the medulla columnar center (Figure 1M’). On the other hand, Fmi–mCherry signal was widely distributed in the R8 axon terminal, including filopodia-like protrusions (Figure 1M”). These protein localization data indicate that Gogo and Fmi functionally cooperate so that R8 axons recognize the center of the medulla column during step 1. At the step 3, the results indicate that Fmi alone promotes vertical filopodia elongation into the M3 layer.

**Figure 1.**
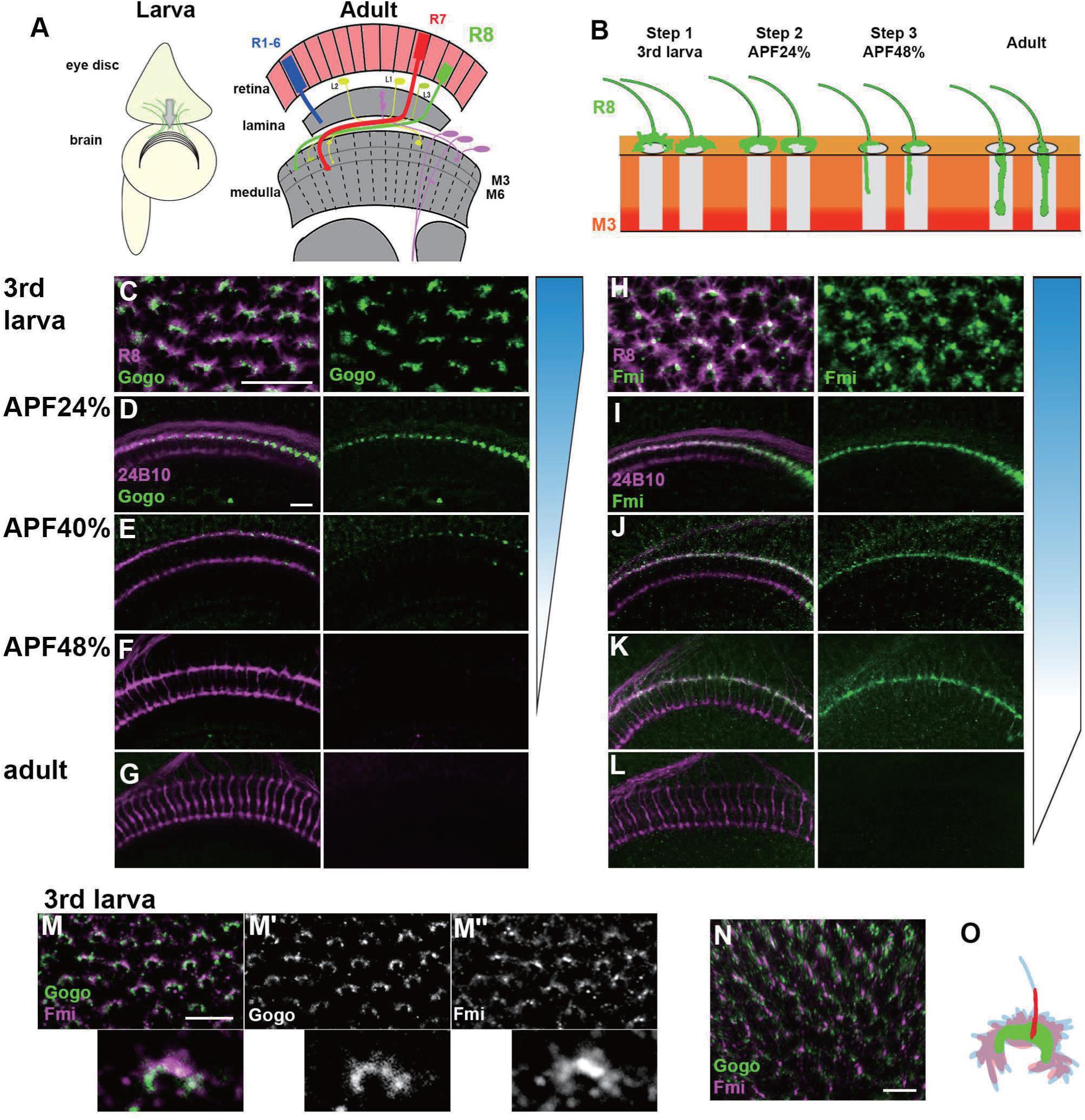
R8 specific labeling of Gogo and Fmi. (A) Schematics of the Drosophila visual system in the third instar larva and the adult. (B) Schematic of the step-wise R8 targeting during development. (C-G) Gogo localization at the terminals of R8 axons (green) visualized by combining Gogo-FsF-GFP and R8 specific FLPase strains (sens-FLP). R8 axons are visualized by myr-RFP (C) or mAb24B10 (D-G) (magenta). (H-L) Fmi protein localization at the terminals of R8 axons (green) visualized by Fmi-FsF-mCherry and R8 specific FLPase strains (sens-FLP). R8 axons are visualized by mCD8GFP (H) or mAb24B10 (I-L) (magenta). (M-O) Localization of Gogo (green) and Fmi (magenta) protein at the tip of R8 axon in 3^rd^ larva (M; 2D, N; 3D). Gogo was strongly enriched at the rim of the horseshoe shaped R8 axon terminal (M’). Fmi was distributed broadly including filopodia (M’’, N). Schematic of Gogo (green) and Fmi (red) expression in R8 cell (blue) (O). Scale bars:10 μm.

### Gogo and Fmi cooperatively guide R8 axons to encircle the columnar center of the medulla

R8 cell-specific null mutant was generated to observe stepwise Gogo and Fmi functions. An RNAi insertion and a heterozygous null mutation were combined (Hakeda-Suzuki et al., 2017), thus resulting in a strong phenotype equivalent to known *gogo* or *fmi* null mutations (Figure S2). In the R8 cell-specific *gogo* mutant, R8 axons correctly targeted each column, but the termini intruded into the medulla columnar center and failed to form a proper horseshoe shape during step 1 (Figures 2A and 2D). Towards step 2, the R8 axonal terminals displayed greater horizontal filopodial extension than normal, thereby enhancing the probability of encountering neighboring *gogo*^-^ R8 axons over time (Figure 2D). This excessive R8 filopodia coincides with the disrupted R8 axon termini lineup and the invasion of layers that are slightly deeper than M1 during steps 2 and 3 (Figures 2B and 2E). Axon bundling and incorrect targeting becomes more prominent later in development. As a result, multiple R8 axons (usually two) were often observed innervating a single column (Figures 2C and 2F). During live imaging, vertical extension could be observed during step 3 in tangled *gogo* R8 axons, thus indicating that it is difficult to uncouple axons that have become tangled (Movies S1 and S2). This clarifies the observation that columnar organization become worse in a larger mutant area compared with a single isolated mutant axon (Tomasi et al., 2008). Furthermore, these data suggest that proper filopodia extension during step 2 is essential for avoiding axon bundling and for promoting a proper array of medulla columns during later development (Figures 2C’, 2F’, and 2G).

**Figure 2.**
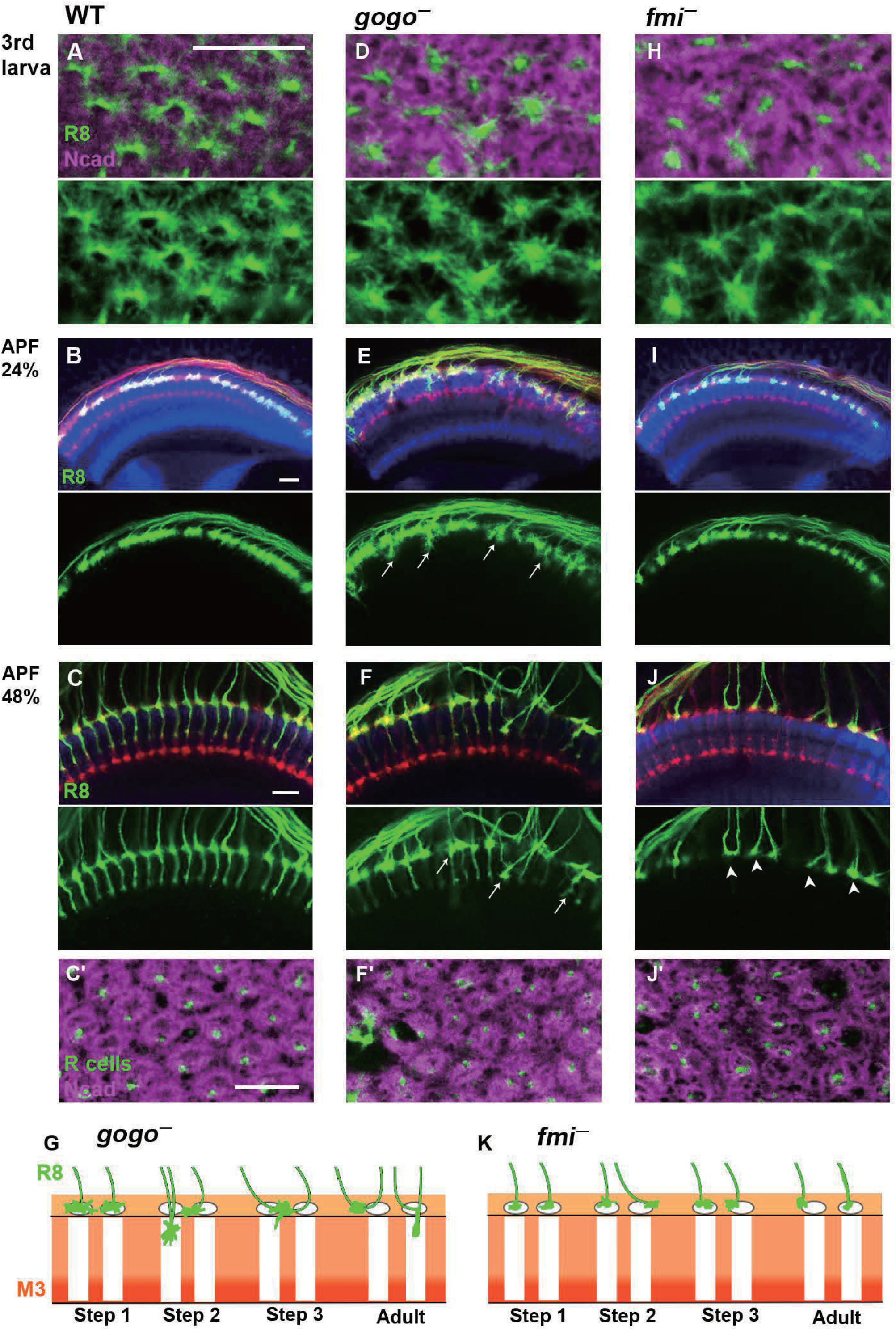
Gogo and Fmi regulates the growth cone dynamics. (A–K) The medulla at the several developmental stages of control (A-C), R8 specific *gogo* mutant (D-G) and R8 specific *fmi* mutant (H-K). R8 axons were labeled with UAS-mCD8GFP (green in A-C, D-F, H-K), R-cell axons with mAb24B10 (red in B, C, E, F, I, J, green in C’, F’, J’), and the medulla layer or column with anti-N-Cadherin (magenta in A, D, H, C’, F’, J’, blue in B, C, E, F, I, J). At APF 24 and 48%, *gogo* mutants show R8 axons bundling phenotype and extend through the R8 temporary layer (E, F arrows), whereas in *fmi* mutants (I, J), R8 axons fail to extend its filopdia vertically towards the M3 layer (arrowheads in J). Medulla columnar pattern labeled with N-Cadherin (magenta) and R axons with mAb24B10 (green) (C’, F’, J’). Schematic of R8 targeting phenotype in *gogo* mutant and *fmi* mutant during development (G and K). Scale bars: 10μm.

Similar to the *gogo* phenotype, the *fmi* mutants had R8 axon terminals that intruded into the medulla columnar center and failed to form a proper horseshoe shape during step 1 (Figure 2H). In contrast to the *gogo* mutant, R8 filopodia horizontal extension was abnormally shortened. As a result, R8 axons maintained their distance from neighboring R8 axons and lined up orderly at the medulla surface during step 2 (Figure 2I). Towards step 3, R8 axons began to lose proper distance among themselves, thus resulting in defective columnar organization (Figures 2C’ and 2J’). We attributed this defects to the initial failure of *fmi* R8 axons to encircle the medulla columnar center during step 1. Moreover, in step 3, *fmi* R8 axons failed to vertically extend their filopodia toward the M3 layer (Figure 2J). These results indicate that Gogo and Fmi function in opposing manners during steps 2 and 3 of R8 axon targeting (Figures 2G, 2K). Given that *gogo* and *fmi* mutants had disorganized medulla columns in later stages (Figure 2J’), it can be concluded that the column center encircling during step 1 is important for R8 axons to follow the correct columnar path and to develop organized arrays.

### Gogo performs a cooperative and antagonistic function toward Fmi

Previous studies that are primarily based on genetic interactions have indicated that Gogo and Fmi must interact to recognize their ligand molecule (Hakeda-Suzuki et al., 2011). Loss-of-function mutations were used to observe any genetic Gogo/Fmi interactions during step 1. The use of RNAi lines to knockdown each gene in an R8-specific manner resulted in morphological defects in the termini of a fraction of R8 axons (38.2% of *gogo*RNAi and 11.9% of *fmi*RNAi; Figures 3A, 3B, and 3D). Double knockdown synergistically enhanced these morphological defects (76.6% of termini; Figures 3C and 3D), thus suggesting that Gogo and Fmi cooperate during step 1 to guide the proper R8 axonal targeting of the columns.

**Figure 3.**
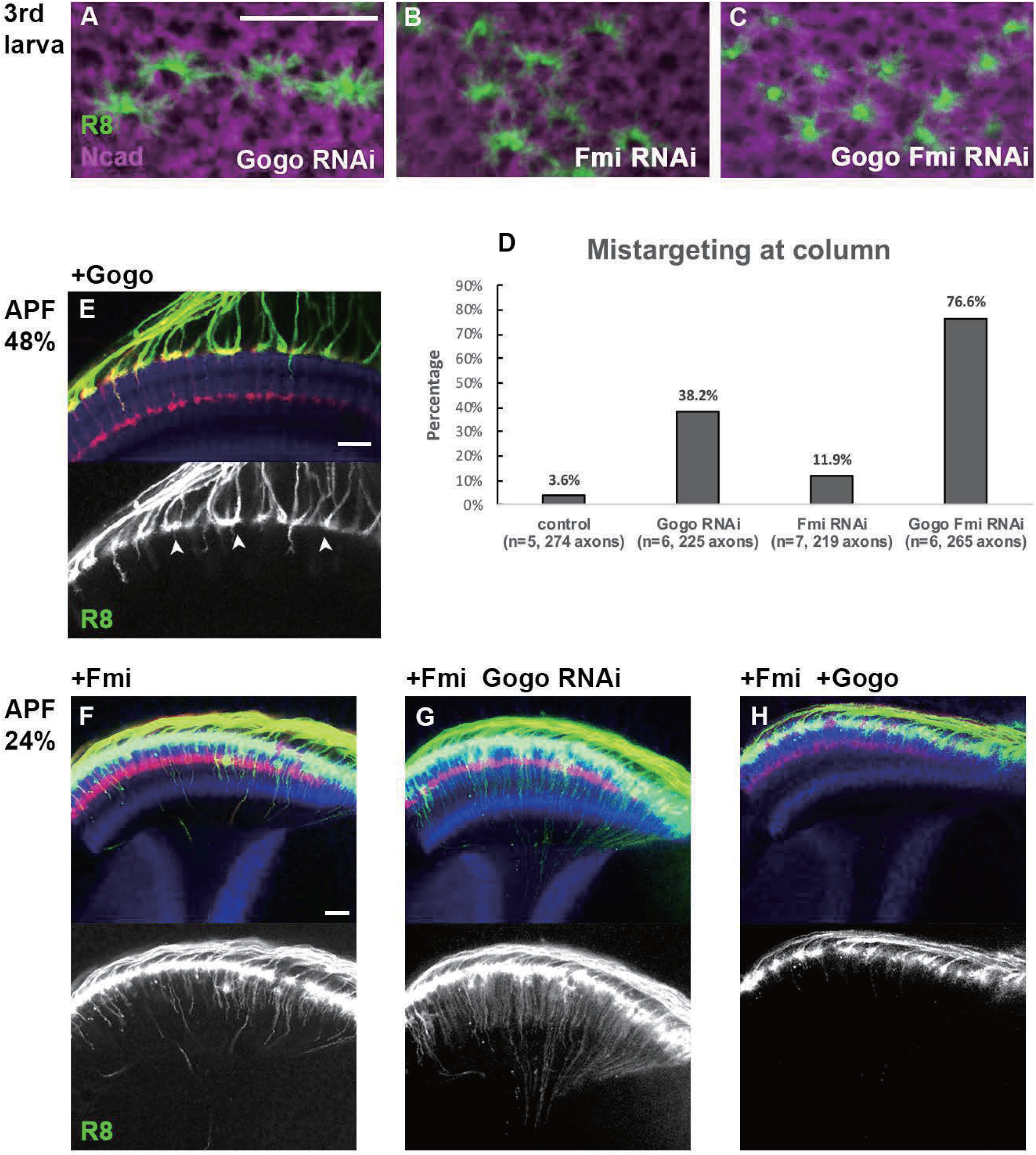
Gogo has dual functions, “cooperative” and “antagonistic” towards Fmi. (A-D) R8 specific knockdown of *gogo* (A)*, fmi* (B), and *gogo*, *fmi* double (C) in R8 specific manner. R8 axons were visualized with UAS-mCD8GFP (green) counterstained with N-Cadherin (magenta). Quantification of the R8 axons mistargeting into the medulla columnar center (D). (E-H) Genetic interaction between *fmi* and *gogo*. R8 axons are labeled with mCD8GFP (green), counterstained with mAb24B10 (red) and N-Cadherin (blue). *gogo*-overexpressed R8 axons fail to extend their filopodia vertically towards the M3 layer (arrowheads in E). In *fmi* overexpression, R8 cells extend their vertical filopodia precociously towards the deeper layer of the medulla at APF 24% (F). The vertical filopodia extension were further promoted by *gogo* RNAi (G) and strongly suppressed by *gogo* overexpression (H). Scale bars: 10μm.

The next set of experiments was attempted to rescue these mutant phenotypes by overexpressing the opposing gene to test whether Gogo and Fmi are mutually compensatory. Fmi overexpression in R8-specific *gogo* mutants did not rescue R8 axon termini morphological defects (Figures S3A and S3C). Likewise, Gogo overexpression in R8-specific *fmi* mutants did not rescue the morphological defects (Figures S3B and S3D). These results indicate that Gogo and Fmi do not have redundant gene functions and cannot compensate for each other.

To investigate the role of Gogo during steps 2 and 3, *gogo* was overexpressed in an R8-specific manner. *Gogo*-overexpressed R8 axons failed to vertically extend their filopodia toward the M3 layer, similar to that in *fmi* mutants (Figure 3E, compared with Figure 2J). Conversely, *fmi*-overexpressed R8 axons vertically extended their filopodia toward the layers much deeper than the wild type and passes through the medulla during step 2 (Figure 3F). To observe the genetic relationship between Gogo and Fmi, Gogo levels were manipulated, and the effect on filopodia extension in *fmi*-overexpressed R8 axons was observed. *gogo* knockdown on an *fmi* overexpression background enhanced premature vertical filopodia extension during step 2 (Figure 3G), thus resulting in the R8 axon bundling phenotype observed at the adult stage (Figures S3E–S3H). Conversely*, gogo* and *fmi* cooverexpression suppressed filopodia extension compared with *fmi* overexpression alone (Figure 3H). These results underscore that Fmi promotes filopodia extension, which is counteracted by Gogo.

Thus, as the development proceeds, Gogo exhibits two-faced attitude towards Fmi, changing its function from cooperation to antagonization.

### The two functions of Gogo are regulated by the same functional ectodomain

To examine how Gogo switches its functional role regarding Fmi, we first checked if Gogo has multiple functional stretches in the extracellular domain that could elicit each function. Gogo has a GOGO domain that contains eight conserved cysteines, a Tsp1 domain, and a CUB domain in its extracellular portion. Previous work has shown that both the GOGO and Tsp1 domains are required for Gogo function (Tomasi et al., 2008). To determine which Gogo ectodomain is required in higher resolution, a smaller segment of each domain was deleted from the genome using CRISPR/Cas9. Severe morphological phenotypes similar to the *gogo* null mutant were observed in any of the small GOGO or Tsp1 domain deletions in step 1 (Figures S4A–S4H). Furthermore, overexpression of Gogo fragment that lacks GOGO or Tsp1 domains, showed weaker filopodia extension suppression in the *fmi* overexpression mutants compared to the full length Gogo overexpression (Figures S4I–S4O). These results demonstrated that GOGO and Tsp1 domains are required in both steps 1 and 2. Therefore, the same stretch of extracellular portion (GOGO–Tsp1) is required for the both functions of Gogo, indicating that switching between two functions of Gogo is not relevant to the extracellular portion during the early developmental stages.

### Gogo localization is dependent on Fmi localization inside filopodia

The functional domain in the extracellular portion of Gogo indicates that Gogo/Fmi interactions occur throughout development, including steps 1 and 2. Previous studies have shown that Gogo and Fmi colocalize at the cell–cell contacts of cultured cells (Hakeda-Suzuki et al., 2011). In order to test it in more *in vivo* situation, we tried to observe the changes of the Gogo or Fmi protein localization at step 1 in the loss- or gain-of-function mutants. Fmi localization was not altered in *gogo* loss-of-function (Figures 4F and 4G) or *gogo* overexpression mutants (Figures 4F and 4H). Conversely, Gogo localization was centrally shifted in the R8 axon termini of *fmi* loss-of-function mutants (Figures 4A and 4B). Under *fmi* overexpression, Gogo localization shifted toward the stalk of the axon terminal, where Fmi accumulates (Figures 4C–4E). Moreover, Gogo localization was shifted along the vertical filopodia stimulated by Fmi to prematurely extend during step 2 (Figures 4I and 4J). These results indicate that Gogo localization is controlled by Fmi. Furthermore, physical Gogo/Fmi interaction controls the formation of the horseshoe structure during step 1 and filopodia extension during step 2.

**Figure 4.**
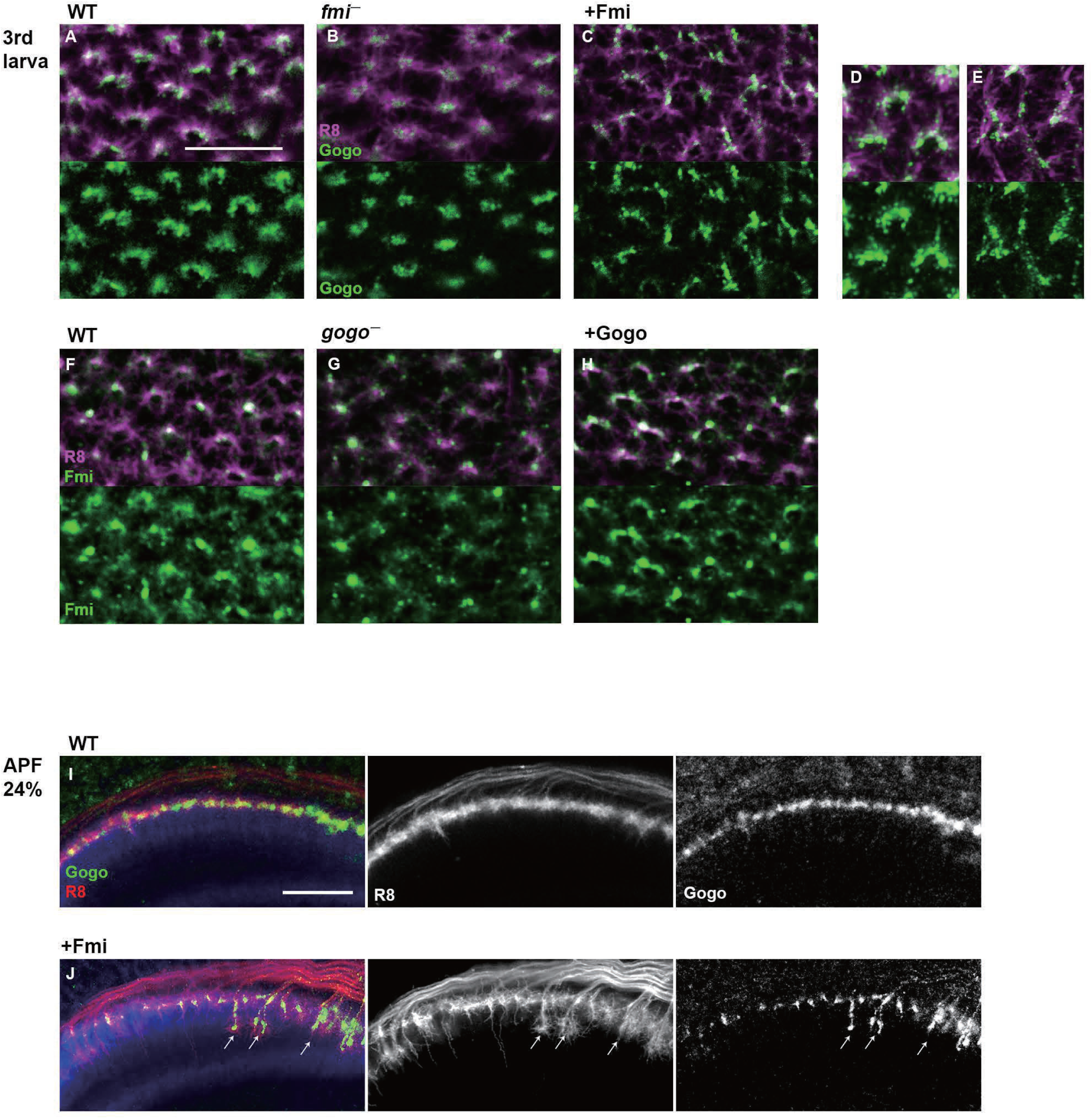
Gogo localization in R8 changes depending on the expression level of Fmi. (A-H) Localization of R8 specific Gogo-GFP and Fmi-mCherry in mutant or overexpression background. R8 axons are labeled with myr-RFP or mCD8GFP. (D-E) 3D images of Gogo localization in R8 cells of wild type (D) or Fmi overexpression (E). In Fmi overexpression, strong Gogo expression is observed at the stalk part of the axon terminal (C and E compared with A and D). Fmi localization did not show notable change in *gogo* mutant (G) nor *gogo* overexpression (H) compared to the wild type (F). (I-J) R8 specific Gogo-GFP (green) at APF24% in wild type (I) and Fmi overexpression (H). R8 axons are labeled with myrRFP (red) and counterstained with N-Cadherin (blue). Gogo protein localizes along the vertical filopodia that prematurely extended at the step 2 (arrows in J compared with I). Scale bars: 10 μm.

### Dephosphorylated and phosphorylated Gogo have distinct functions toward Fmi

We next tested whether cytoplasmic domain of Gogo serves as a switch to change between its two-faced functions. Previous studies suggest that the cytoplasmic domain of Gogo is important for Gogo/Fmi collaborative functions, while they interact *in cis* (Hakeda-Suzuki et al., 2011; Tomasi et al., 2008). It has also been shown that the YYD tripeptide motif in the cytoplasmic domain is required for Gogo function (Mann et al., 2012). Furthermore, Tyr1019 and Tyr1020 are known as the true phosphorylation sites *in vivo* (Mann et al., 2012). To test whether Gogo phosphorylation regulation is required during step 1, the Gogo phosphomimetic form (GogoDDD), nonphosphomimetic form (GogoFFD) and deletion of the entire cytoplasmic domain (GogoΔC) were used to rescue the *gogo* mutant phenotype. GogoDDD and GogoΔC were unable to rescue the mutant morphological phenotype, whereas wild-type Gogo and GogoFFD significantly rescued the phenotype during step 1 (Figures 5A–5F). These results indicate that the unphosphorylated YYD motif of the cytoplasmic domain is required for R8 axons to correctly recognize the medulla column and encircle the columnar center.

**Figure 5.**
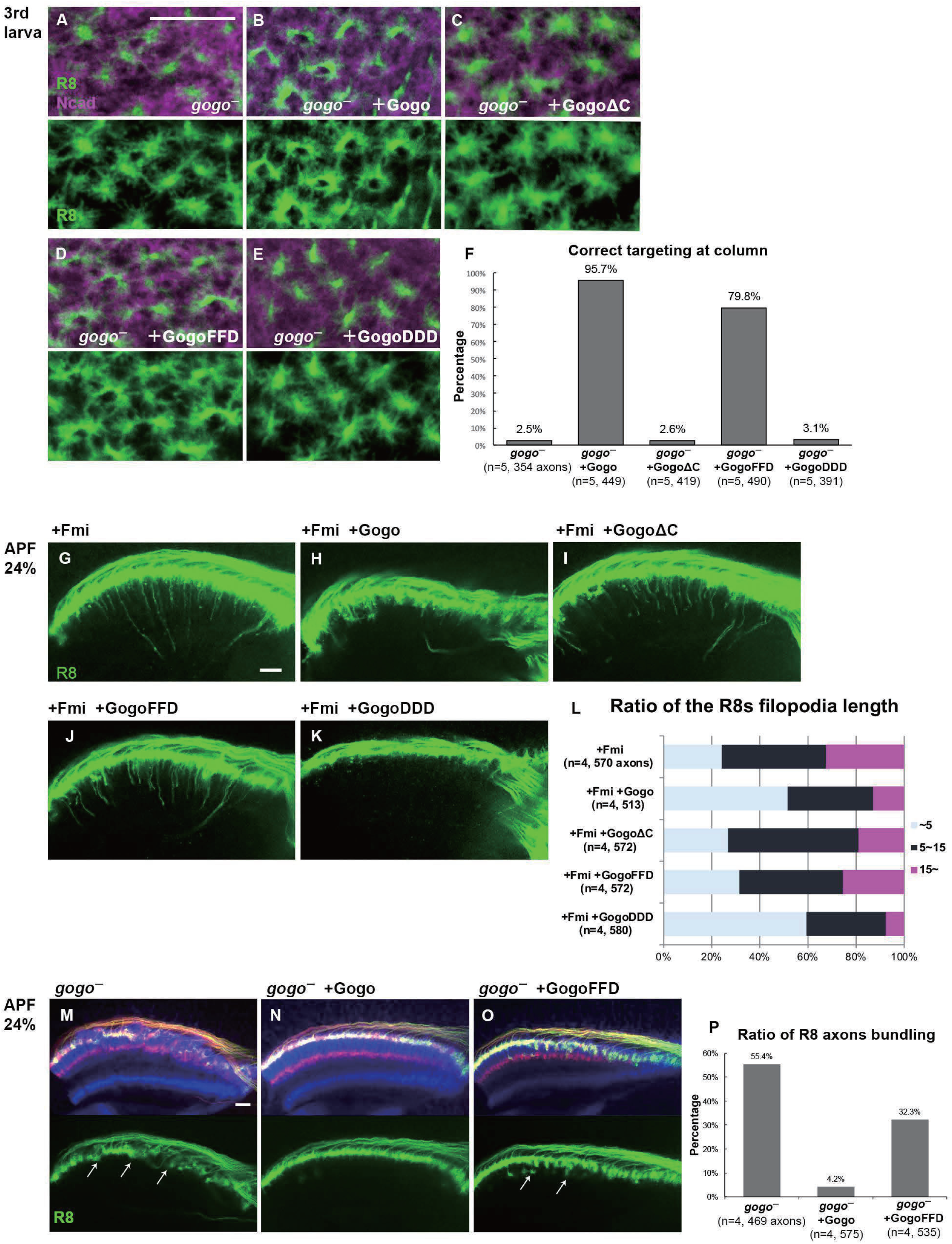
Dual function of Gogo regulated by the phosphorylation of YYD motif. (A-F) *gogo* rescue experiments in a background of *gogo*[H1675] / *gogo*[D1600]. R8 axons are visualized with mCD8GFP (green), and column are labeled with N-Cadherin (magenta). The targeting defects of *gogo* mutants (A) are almost completely rescued with wild-type Gogo (B) and GogoFFD (D: non-phospho-mimetic) but were not rescued by GogoΔC (C) or GogoDDD (E: phospho-mimetic). (F) Quantification of R8 axons targeting with correct column position. (G-L) Horizontal images of R8 cells that expresses GogoFFD or GogoDDD in Fmi overexpression background at APF24%. R8 filopodia elongation was significantly repressed by wild-type Gogo (H) or GogoDDD (K), but not with GogoΔC (I) nor GogoFFD (J). Quantification of R8 axons filopodia length (L). The longest filopodia in a 3D image was measured from each axon. The length of filopodia is divided into 3 classes of ∼ 5μm (Light blue), 5-15 m (dark blue), 15 μm ∼(magenta). (M-P) Ectopic filopodia extension and axon bundling (arrows in M) in *gogo* mutant (*gogo*[H1675] / *gogo*[D1600]) was rescued by Gogo (N) but not by GogoFFD (arrows in O) at APF 24%. Quantification of the bundling R8 axons (P). Scale bars: 10 μm.

Next, we sought to determine which Gogo form is functional during filopodia extension in step 2. The GogoFFD and GogoDDD transgenes were inserted in *fmi*-overexpressed flies (Figures 5G–5L). GogoFFD did not suppress filopodia extension (Figures 5J and 5L), but GogoDDD did (Figures 5K and 5L). This indicates that the phosphorylated form of Gogo is required for filopodia suppression.

In previous studies, GogoFFD rescued the R axon targeting defects in adult stage to a considerable extent (Mann et al., 2012). However, in the current study at earlier stages, GogoFFD did not completely rescue ectopic filopodia extension and axon bundling, thus resulting in a slightly premature R8 termini intrusion into the medulla neuropil during step 2 (Figures 5M–5P). Therefore, Gogo phosphorylation must occur sometime between steps 1 and 2 to suppress excessive filopodia formation and extension during normal R8 axon development. These results suggest that Gogo phosphorylation statuses have distinct functions in axon pathfinding to form complex functional neuronal circuit.

### Suppression of Fmi by phosphorylated Gogo is mediated via adducin

Gogo interacts with the actin-capping protein Hu-li tai shao (Hts, *Drosophila* adducing homolog) to control R8 neuron axonal extension (Ohler et al., 2011). Thus, we hypothesized that the Gogo function that suppresses filopodia relies on the actin-capping ability of Hts. Thereafter, R8-specific *hts*^—^ mutant was analyzed. During step 2, *hts*^—^ R8 axon termini had excessive and random filopodia extensions and an axon–axon bundling phenotype similar to *gogo*^—^ mutants (Figure S5A), thus suggesting that Hts works with Gogo to prevent excessive filopodia extension. To determine which Gogo form works with Hts, Hts was cooverexpressed with GogoDDD or GogoFFD, and this was observed during step 3 (Figure S5B) and in adulthood (Figure S5C). Wild-type Gogo or GogoDDD overexpression partially suppressed filopodia extension (Figure S5B). GogoDDD*/*Hts coexpression, but not GogoFFD*/*Hts coexpression, synergistically suppressed filopodia extension or resulted in R8 axon stalling at the medulla surface layers (Figures S5C and S5D). These data indicate that phosphorylated Gogo sends signals via Hts to suppress filopodia extension.

### Glial cell insulin signal is critical for Gogo phosphorylation

The Gogo/Fmi interaction phenotype can be considered “cooperative” in step 1 but changes to “antagonistic” in step 2 (Figures 2 and 3). This indicates that Gogo is phosphorylated during the transition from steps1 to 2, but the mechanism is unclear. Previous work indicates that DInR phosphorylates Gogo and is important for its function (Mann et al., 2012). DInR has tyrosine kinase activity and is known to phosphorylate the YYD motif. Therefore, R8-specific *dinr*^—^ mutant was created. During step 2, the *dinr*^—^ mutant R8 axons displayed a similar phenotype to the GogoFFD rescue and exhibited random filopodia extensions, thus resulting in R8 axon bundling and the premature invasion of the deeper medulla layers (Figure 6A, compared with Figure 5O).

**Figure 6.**
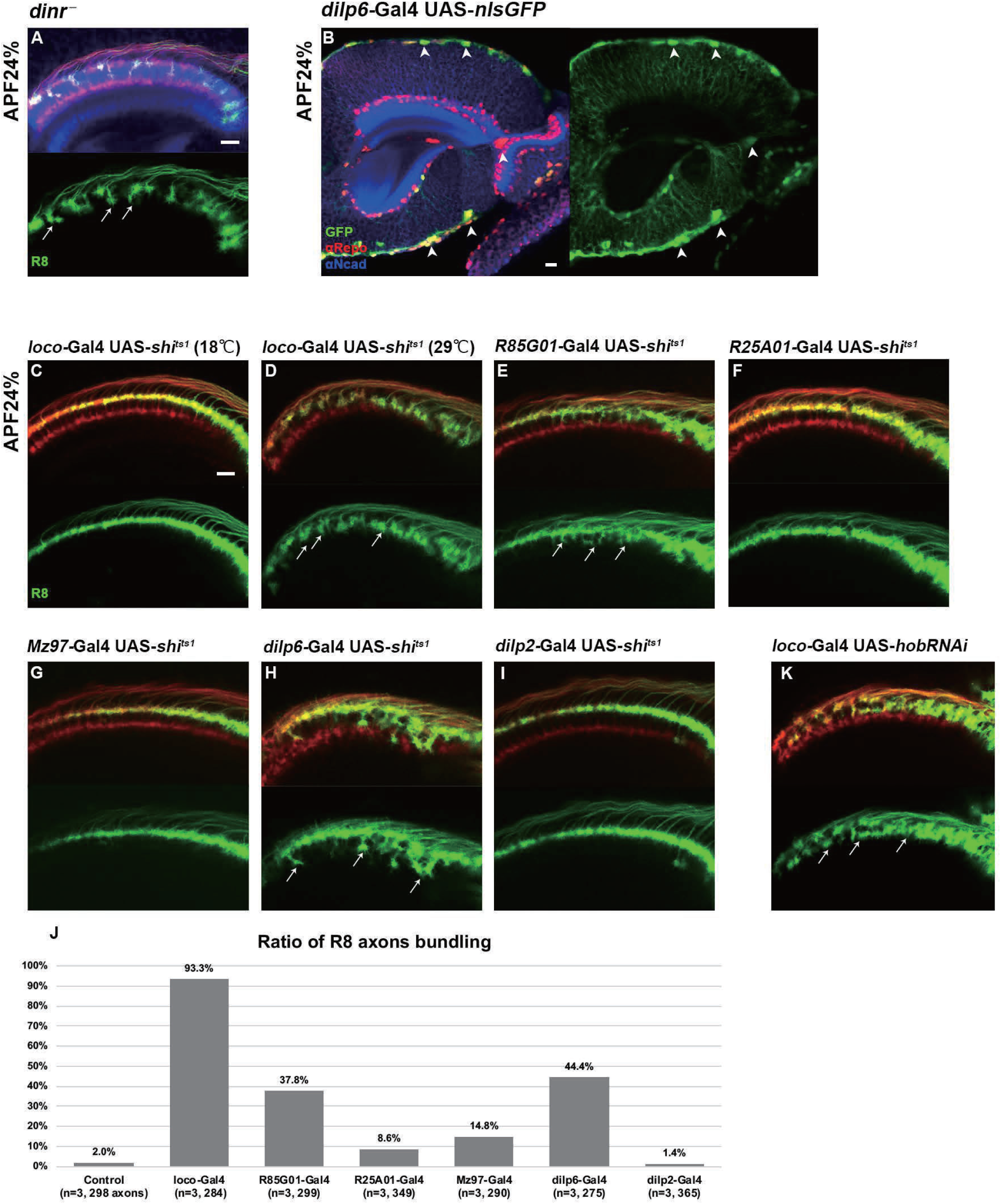
Glial insulin switches the Gogo-Fmi function from “cooperative” to “antagonistic”. (A) R8 specific *dinr* mutant at APF 24%. R8 axons were labeled with mCD8GFP (green) counterstained with mAb24B10 (red) and N-Cadherin (blue). R8 axons bundle to each other, and precociously invade into deeper medulla layers (arrows). (B) Dilp6-Gal4 expression monitored by nuclear GFP reporter (green) is seen mainly in cortex and surface glial cells in optic lobe at APF 24% (arrowheads). Glial cells are labeled with anti-repo (red), optic neuropil with anti-N-Cadherin (blue). (C-J) The secretion of the Dilp are blocked by UAS-*shi^ts1^* in all glia (*loco-Gal4,* C and D), surface and cortex glia (GMR85G01-Gal4, E), wrapping and neuropil glia (GMR25A01-Gal4, F or Mz97-Gal4, G), *dilp6*-Gal4 (H) or *dilp2*-Gal4 (I). At APF 24%, R8 axons labeled by myr-tdTomato (green) showed bundling phenotype in surface and cortex glia specific *shi^ts1^* expression (arrows in D, E and H). Quantification of the bundling R8 axons (J). (K) Glia-specific inhibition of Dilp secretion caused by *hobbit* RNAi expressed under *loco*-Gal4 driver. R8 axons bundled to each other, and precociously invade the deeper medulla layers (arrows). Scale bars: 10 μm.

The next question was to determine how DInRs on R8 axons receive insulin signals. Previous gene expression studies in the developing optic lobe revealed that among the eight *dilp* genes, *dilp6* is expressed in glia cells in *Drosophila* (Fernandes et al., 2017; Okamoto and Nishimura, 2015; Rossi and Fernandes, 2018; Sousa-Nunes et al., 2011). By using Gal4 lines, *dilp6* expression in the surface and cortex glia was observed at all developmental stages (Figures 6B and S6A–S6I). To identify whether glia contributes to Gogo phosphorylation in R8 axons, glial-specific protein secretion was blocked during step 2. Dynamin is known to control peptide secretion, including insulin-like peptides (Wong et al., 2015). The temperature-sensitive dynamin mutant (*shibire^ts1^* [*shi^ts1^*]) was specifically expressed in glial cells to block dilp secretion. This produced a defective phenotype similar to the *dinr* mutants; R8 axons showed random filopodia extensions and bundling with premature invasion into deeper medulla layers (Figures 6C and 6D). These defects were also observed when *shi^ts1^* was specifically overexpressed in surface and cortex glia cells (Figures 6E, 6H, and 6J). Conversely, we could not see any defects when we block the protein secretion from insulin producing cells (IPC) (Figures 6I and 6J) or other types of glia cells, including medulla neuropil glia and Chiasm glia (Figures 6F, 6G and 6J).

The *hobbit* gene is known to regulate Dilp secretion (Neuman and Bashirullah, 2018). Therefore, *hobbit* was knocked down to block Dilp secretion specifically in glial cells. This produced a similar phenotype as the *dinr* mutant, thus supporting the idea that glial Dilp controls R8 axonal targeting (Figure 6K). Unfortunately, R8 axonal targeting defects could not be detected in *dilp6* knockdown mutants probably because the dilps have redundant functions (Figure S6). Although the precise dilp that regulates this process could not be identified, these results suggest that R8 neuron DInR phosphorylates the Gogo cytoplasmic YYD motif upon receiving glia-derived insulin signals during step 2.

### Glia supplies Fmi that interacts with R8 axons in the columnar center

We have shown that Gogo and Fmi direct R8 axons to recognize the columnar center. However, the component that R8 recognizes during step 1 is unclear. We hypothesized that the Fmi located on R8 axons functions as a cadherin and homophilically adheres with Fmi on neighboring cells, thereby allowing R8 axons to correctly target the medulla. R7, R8, and Mi1 neurons are known to be the core members during the earliest medulla column formation step (Trush et al., 2019). To test whether functional Fmi is located on R7 or Mi1, Fmi was specifically knocked down in R7 or Mi1 neurons. This did not result in detectable defects in the overall R8 axon targeting or termini morphology (Figures S7A and S7B). During the analysis of glial cell function for insulin signaling, we noticed a firm localization of the Fmi protein at the glial protrusion in the columnar center at the step 1 (Figures 7A and 7B). Considering that glial cells also contact R8 axons, glia-specific *fmi* mutants were created. The phenotypes were similar to that of the *gogo* and *fmi* R8 mutants (Figures 2D and 2H). R8 axon termini in the optic lobe of these mutants failed to encircle the columnar center and intrude into the central area (Figures 7C and 7D). In the step 3, columnar organization was disturbed in glia-specific *fmi* mutants. Proper distance was not maintained between R8 axons and the fine columnar array was disrupted (Figures 7C’ and 7D’).

**Figure 7.**
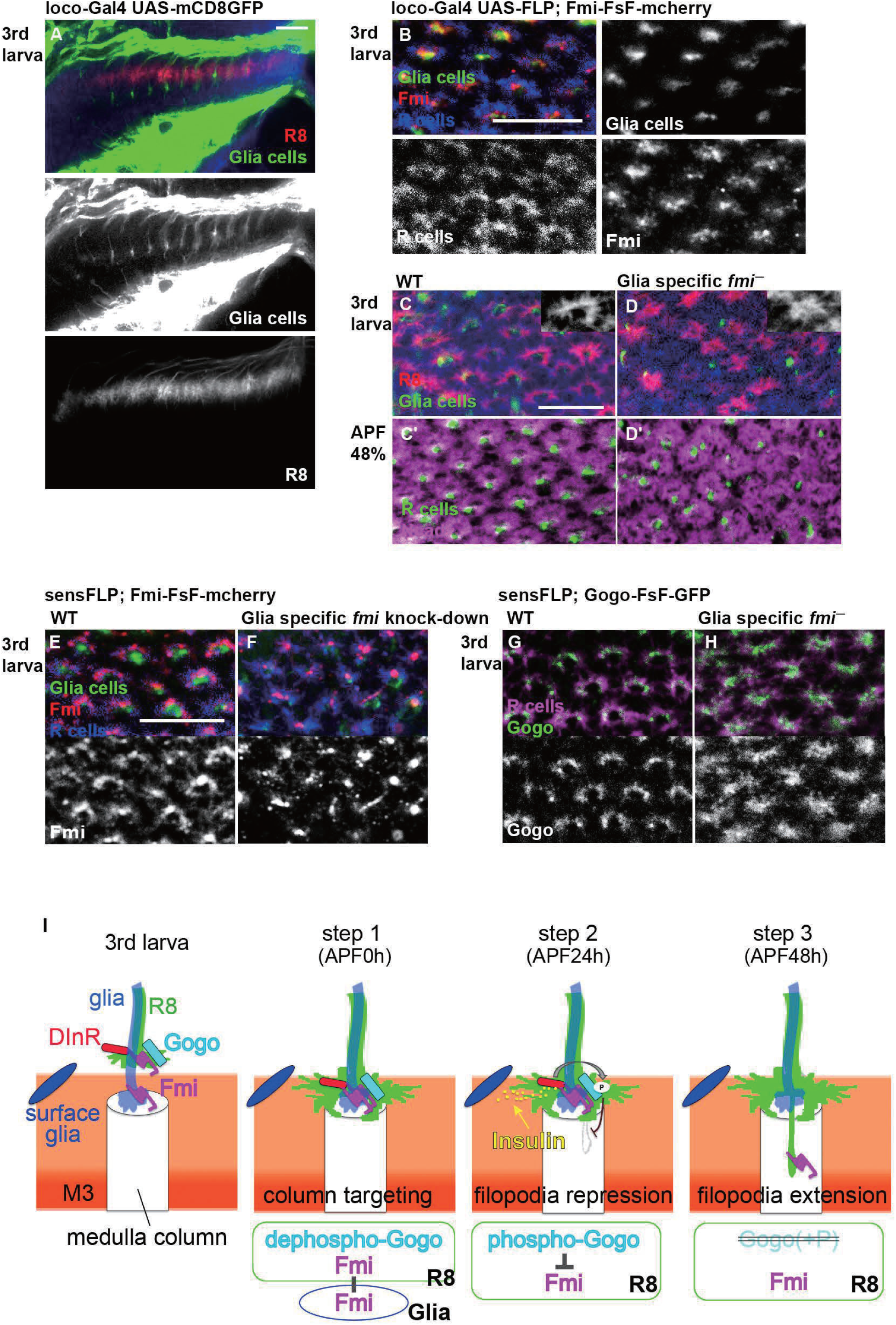
Glial Fmi and R8 Gogo/Fmi instruct R8 to recognize the columnar center. (A) R8 terminals visualized with myr-tdTomato (red, white) and Glial cells visualized with mCD8GFP (green) counterstained with N-Cadherin (blue). (B) The localization of Fmi (red) in the Glial cells (green; mCD8GFP) of the medulla neuropil counterstained with mAb24B10 (blue). (C, D) Wild type (C) and glial-specific *fmi* mutant (D) medulla in 3^rd^ larva. Labeling is same as in (A) (C’, D’) glial-specific *fmi* mutant medulla at APF 48%. Medulla columnar pattern is labeled with N-Cadherin (magenta) and R axons with mAb24B10 (green). (E, F) The localization of Glial cells (green) in Medulla neuropil and Fmi-mCherry (red) in R8 cells. R axons are labeled with mAb24B10 (blue). (G, H) Localization of R8 specific Gogo-GFP (green) in glia specific *fmi* mutant. R axons are labeled with mAb24B10 (magenta). Scale bars: 10μ (I) Model for the interaction between two-faced Gogo and Fmi to navigate R8 axons. In the step 1, non-phosphorylated Gogo/Fmi at R8 termini interact *in trans* with Fmi which is localized on the glial surface, and collaborate to correctly recognize the medulla columnar center. In the step 2, Gogo is phosphorylated by the Insulin signal derived from surface & cortex glia. Phospho-Gogo antagonizes Fmi, thereby suppresses filopodia extention. In the step 3, Fmi alone brings R8 axon to the M3 layer, since Gogo protein no longer exists in R8 axons.

Changes in R8 axon Gogo and Fmi localization were observed in glia-specific *fmi* mutants to further assess the functional relationship between glial Fmi and R8 Gogo/Fmi. R8 axon Fmi localization was weaker in the filopodia tips and accumulated in the axon termini stalk (Figures 7E and 7F). R8 axon Gogo localization was more diffuse throughout the entire termini structure, including in filopodia (Figures 7G and 7H). These localization changes indicate that Gogo and Fmi relocate from the R8 axon horseshoe rim to other regions when R8 axon Fmi cannot bind to glial Fmi. These results also indicate that the *in trans* interaction between glial Fmi and R8 Gogo/Fmi mediates precise R8 axon recognition with the medulla columnar center, including the formation of a horseshoe structure. This interaction is mediated by nonphosphorylated Gogo, and later the phosphorylation of Gogo switches the Gogo/Fmi function from “collaborative” to “antagonistic” (Figure 5).

These results suggest that the glial insulin signal controls the phosphorylation status of Gogo, which regulates the growth cone dynamics of R8 and mediates axon–glia and axon–axon interactions (Figure 6). This mechanism maintains a consistent distance between R8 axons, enables ordered R8 targeting of the column, and eventually contributes to the formation of the organized array of the medulla columns (Figure 7I).

### Gogo and Fmi function in mushroom body

Gogo and Fmi are broadly expressed in nervous systems outside of the visual system. Previous work has shown that Gogo and Fmi function in dendrite formation during the embryonic stage (Hakeda and Suzuki, 2013; Hakeda-Suzuki et al., 2011). The current experiment demonstrates that Gogo functionally regulates correct axon targeting in the mushroom body similar to the visual system (Figure 8). The mushroom body is a higher center for olfactory learning and memory (de Belle and Heisenberg, 1994). Previous studies have shown that *fmi* mutant axons also have targeting defects in mushroom body neurons (Reuter et al., 2003). A side-by-side analysis of *gogo* mutants and *fmi* knockdown mutants revealed that the phenotypes were similar but not identical (Figure 8I). When *gogo* was knocked down in *fmi* overexpression mutants, there was more postaggregation or branch guidance defects than mere *gogo* single knockdown (Figure 8J). These phenotypic patterns appear to be similar to the *gogo* mutants. This result demonstrates that *fmi* overexpression can enhance the *gogo* knockdown phenotype, such as R8 axon bundling. In the mushroom body, α/β kenyon cells extend their axons to follow α’/β’, whereas α’/β’ cells follow γ lobe axons (Billuart et al., 2001; Lee et al., 1999; Reuter et al., 2003) (Figure 8A). Similarly, R8 axons extend to follow glial cell protrusions in the visual system. These findings indicate that Gogo/Fmi interactions are broadly utilized in the *Drosophila* nervous system. Given that Fmi is broadly functionally conserved among species (Berger-Muller and Suzuki, 2011; Rapti et al., 2017; Shi et al., 2014; Tissir et al., 2002), elucidating the conserved function of Gogo/Fmi interactions in the *Drosophila* brain can provide valuable insights into the formation of higher-order nervous systems, such as the mammalian brain.

**Figure 8.**
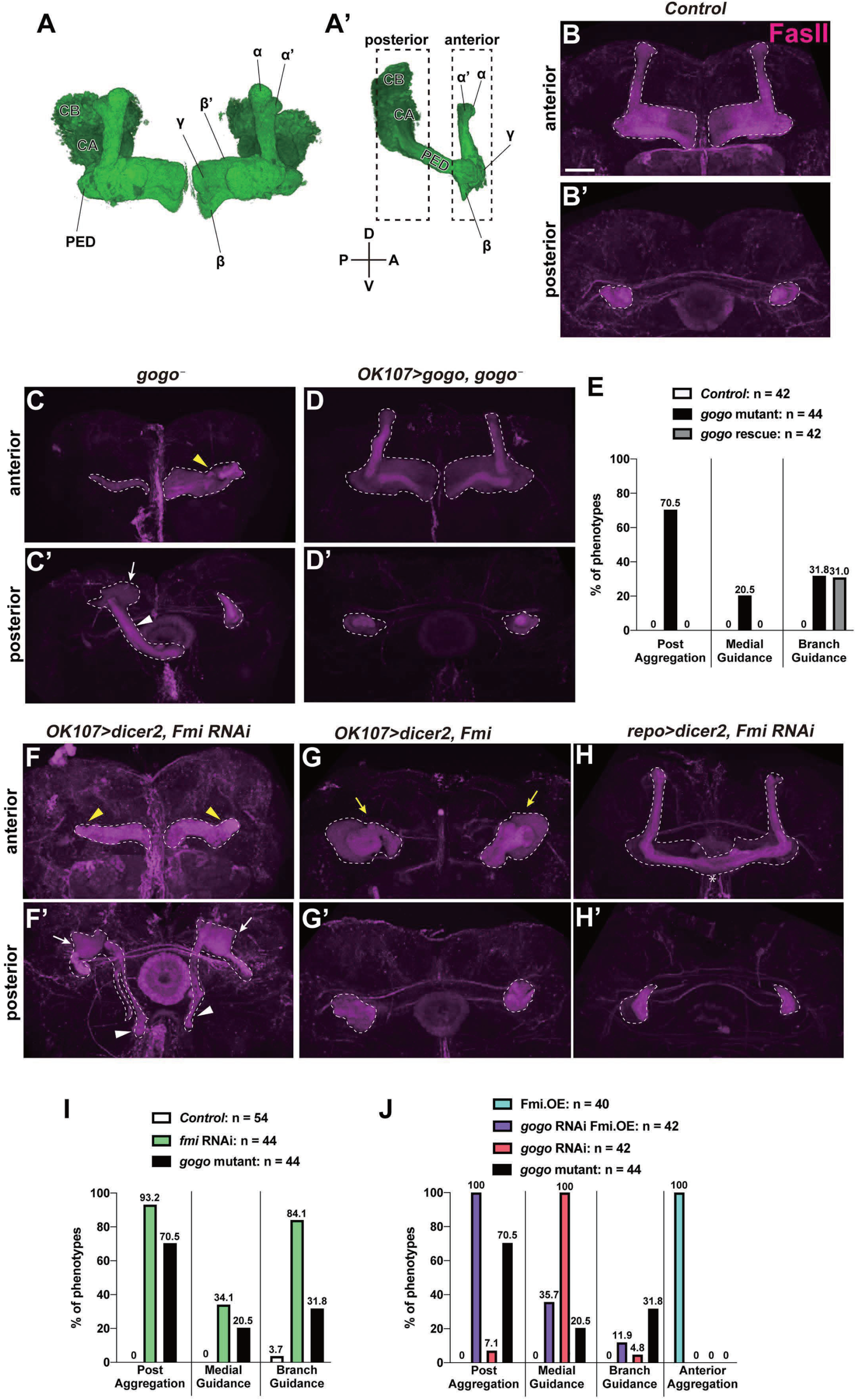
Genetic interaction between *gogo* and *fmi* in mushroom body. (A-A’) Structure of mushroom bodies (MBs) in adult. PED, CA, and CB denote peduncle, calyx, and cell bodies, respectively. Dashed line rectangles in (A’) indicate the anterior region and posterior region in following figures. (B-E) Representative images showing the MBs of the control (B), *gogo* mutant (C), and *gogo* mutant expressing the MB-specific full-length *gogo* (D), γ and α/β lobes were visualized by anti-FasII (magenta). The dashed line in (B-D) demarcates MBs. *gogo* mutant flies displayed axonal branch guidance defects of α/β lobes (yellow arrowhead in C), lobe aggregation in posterior side (white arrow in C’), and misguidance towards the medial side directly from calyx (white arrowhead in C’). (E) Quantification of the rescue experiments of the *gogo* mutant phenotypes. (F-H) Representative images of MBs of *OK107-Gal4*x*UAS-Fmi^RNAi^* (F), *OK107-Gal4*x*UAS-Fmi* (G), and *repo-Gal4*x*UAS-Fmi^RNAi^* (H). γ and α/β lobes were visualized by anti-FasII (magenta). The dashed line demarcates MBs. MB-specific knockdown of Fmi displayed branch guidance defects (yellow arrowhead in F), posterior aggregation (white arrow in F’), and misguidance towards the medial side (white arrowhead in F’). MB-specific overexpression of Fmi caused the lobe aggregation in anterior side (yellow arrow in G). Glial-specific knockdown of *fmi* displayed only the extension of β lobes (asterisk in H; n = 30). (I) Quantification of the MB lobe phenotypes of *fmi* knockdown. (J) Quantification of the lobe phenotypes of genetic interactions between *gogo* and *fmi*. Scale bar: 30 µm.

## Discussion

The current study demonstrated that R8 axons are guided in a stepwise manner via Gogo/Fmi interactions that initially have a collaborative relationship that switches to antagonistic during the development of the visual system (Figure 7I). During step 1, dephosphorylated Gogo interacts with Fmi in cis and cooperatively functions to navigate R8 axons to the correct target. During this stage, R8 Gogo interacts with glial Fmi to locate the column center and enable R8 axon terminals to form a horseshoe-like morphology that encircle the central area of the medulla column. During step 2, Gogo is phosphorylated by the insulin signal derived from the surface and cortex glia. Phosphorylated Gogo antagonizes Fmi via Hts (adducin) to suppress filopodia extension. During step 3, Gogo is no longer expressed in R8 axons; therefore, Fmi alone navigates R8 axons to the M3 layer. Two Gogo states control axon–axon interaction to maintain R8 axon distance and axon–column interaction for proper column targeting.

### Gogo and Fmi cooperatively mediate R8 axon–column interaction during step 1

During step 1, R8 axon terminals form a horseshoe-like shape and encircle the medulla column center. In this step, Gogo and Fmi protein localize at the R8 axon terminal fringe surrounding the medulla center and appear to interact *in cis* (Figures 1M). Because GogoFFD rescued the *gogo* mutant phenotype at this time point, it can be deduced that only the non-phosphorylated version is required. (Figures 5D–5F).

We asked what does phosphorylation do to the function of Gogo. Gogo/Fmi interactions *in cis* occur with the same affinity regardless of the Gogo phosphomimetic version in S2 cultured cells (Mann et al., 2012). Furthermore, GogoDDD and GogoFFD localization did not differ in the R8 axon termini during step 1 *in vivo* (Figure S5E), thus suggesting that the phosphorylation status of Gogo does not change the molecular affinity of Gogo/Fmi.

Gogo phosphorylation may control multiple aspects of this process, including downstream Gogo/Fmi intracellular signaling. The Fmi downstream signaling pathway components that regulate dendrite formation or planar cell polarity (PCP) are well known (Berger-Muller and Suzuki, 2011; Kimura et al., 2006; Li et al., 2016; Lu et al., 1999; Usui et al., 1999; Wang et al., 2016). Previous studies have shown that PCP complex mutants display normal R8 axon targeting in adulthood (Hakeda-Suzuki et al., 2011). Moreover, the RNAi knockdown of components that are thought to regulate the dendrite formation downstream of Fmi, such as PCP complexes and G alpha proteins, did not result in defective R8 axon targeting phenotypes (data not shown). Functionally, the deletion of the intracellular domain of Fmi can promote filopodia elongation but does not mediate column center encircling (Figures S5F and S5G). Given that the Gogo cytoplasmic domain is also required for column center encircling (Figure 5C), the Gogo/Fmi interaction in step 1 may send signals via the Gogo and Fmi cytoplasmic domains.

Previous studies have reported that Gogo/Fmi cooverexpression in R7 axons redirects them to the M3 layer. This occurs when GogoFFD, but not GogoDDD, is expressed (Mann et al., 2012). The observation of this redirection process during development showed that R7 axons do not extend in a stepwise manner such as R8 axons but retreat to the M3 layer from M6 (Figures S7C and S7D). This indicates that Gogo/Fmi cooverexpression does not form a code for M3 targeting but promotes cytoskeletal reorganization, which might lead to R7 axon retraction. The adhesion to the M3 Fmi *in trans*, will be an ectopic situation. Consistent with this idea, R7 retraction was recapitulated by overexpressing Rho by using GMR–Rho1 (Figure S7). It is well known that Rho promotes cytoskeletal reorganization by activating caspase (Aznar and Lacal, 2001; Barrett et al., 1997; Mashima et al., 1999; Shi and Wei, 2007; Sokolowski et al., 2014). The retraction ratio was also enhanced by cooverexpressing Gogo (Figures S7H and S7J).

Strong Gogo/Fmi cooverexpression results in serious cell death in the retina (Tomasi et al., 2008), with greater cell death in GogoFFD than in GogoDDD. If these cell deaths are the result of increased cytoskeleton reorganization, it may indicate that GogoFFD and Fmi cooperatively regulate the cytoskeleton in various steps throughout photoreceptor development. These growth cone dynamics resemble each other when R8 axon Gogo/Fmi interact with glial Fmi to form the horseshoe shape via cytoskeletal reorganization during step 1 (Figures 2 and 7). However, the manner in which GogoFFD sends signals via downstream components and regulates cytoskeleton reorganization is unknown; this must be addressed in the future.

### Glia interact with R8 cells to guide R8 axons during step 1

This study shows that Gogo/Fmi at the R8 termini interacts in trans with Fmi, which is localized on the glial surface during step 1 (Figure 7). Related to these findings, N-cadherin (Ncad) plays a role in medulla column formation (Trush et al., 2019). *Ncad* mutant R8 axons has a defect in targeting the medulla column, which is thought to be due to the difference in adhesive properties of the axons in the column, i.e., the differential adhesion hypothesis (DAH) (Foty and Steinberg, 2005; Murakawa and Togashi, 2015; Trush et al., 2019). In this system, axons with greater Ncad expression tend to target the center of the column, whereas those with lower expression tend to surround the edge of the column border. Ncad overexpression in R8 axons results in changes in termini morphology and in the coverage of the entire medulla column surface (Trush et al., 2019).

In the current studies, Fmi overexpression in the R8 axon termini did not change the horseshoe shape (Figure S3A). However, *fmi* mutant in R8 axons resulted in misguided filopodia invading the column center; this does not support the DAH theory for Fmi (Figure 2C). Therefore, we suggest that as a cadherin, Fmi interacts homophilically in trans as Fmi/Fmi between glia and R8 cells. Conversely, Gogo interacts with Fmi in cis to form Gogo/Fmi on the R8 membrane. Distinct signaling regulation via Gogo and Fmi cytoplasmic domains enables R8 axons to correctly target the medulla column.

One interesting observation is that Gogo localization differed between R8 axon- and glia-specific *fmi* mutants: Gogo protein localization is more diffuse in R8 *fmi* mutant than in glial *fmi* mutants (Figures 4B and 7H). It is known that Gogo and Fmi do not interact *in trans*, which was shown in cell culture systems (Hakeda-Suzuki et al., 2011). These observations suggest that Gogo/Fmi is not only interacting with glial Fmi, but the Gogo ligand (factor X) exists on the glial membrane and interacts with Gogo as Gogo/factor X, in addition to the Fmi/Fmi interaction. The functional role of factor X on glial cells is unknown. Therefore, it is important to identify the role of factor X to reveal the functional significance of glial-derived signaling during step 1 of R8 axon targeting.

### Temporal and spatial regulation of Gogo phosphorylation status by glia

In step 1, R8 axons interact with Fmi on glia cells. In step 2, R8 axons receive insulin from surface and cortex glia. However, insulin expression started at the transcriptional level during step 1 (Figures S6H and S6I); therefore, the temporal relationship of Gogo phosphorylation regulated by insulin is unclear.

One scenario is that it is regulated via changes in the relative position between the glia and medulla during development. Glia position changes across steps 1 to 2 as the entire brain structure changes. There is a huge distance between glia and the medulla neuropil during step 1 that drastically shrinks by step 2. This physical distance between glia and R8 axon termini might influence the reception efficiency of insulin.

The second scenario is that there might be a slow transition between the nonphosphorylated state to the phosphorylated state. Gogo coexists as two phosphorylated states in the tip of R8 axons when R8 axons reach the medulla column. Only the microlocalization of the two phosphorylated states might be differently regulated. The shape of the growth cone was shown to be different between GogoFFD rescue and wild-type rescue in the *gogo* mutant during step 1 (Figures 5B and 5D). This difference might be due to Gogo phosphorylation and may occur even in wild-type overexpression mutants that gained the ability to suppress filopodia extension in step 1.

The transition of total Gogo protein levels in the R8 axons also appeared to be slow. This is based on the observation that *gogo*–Gal4 strain, in which Gal4 is knocked into the *gogo* intron locus by using the MiMIC system (Venken et al., 2011), loses GFP protein levels (monitored by UAS-mCD8GFP, Figure S1) gradually, similar to the gradual decrease of Gogo–GFP fusion protein during the midpupal stages. This indicates that Gogo protein perdurance is similar to mCD8GFP and is not actively degraded by the ubiquitin–proteasome pathway. In summary, the stepwise regulation of R8 axon extension occurs in a precise temporal manner even within developmental steps. However, it is not clear how this is regulated because the transition of Gogo phosphorylation and the decrease in protein level are slow.

### Gogo acts antagonistically against Fmi in R8 axon–axon interactions during step 2

Filopodia are formed by actin polymerization. If the concentration is above a specific threshold, in vitro experiments suggest that actin can polymerize itself. The actin concentration *in vivo* is typically higher than the threshold. This suggests that actin should primarily be controlled by factors that interfere with or suppress uncoordinated actin fiber polymerization in the R8 axon growth cone (Pantaloni et al., 2001; Pollard and Borisy, 2003).

To prevent filopodia extension, actin-capping proteins bind to the end of F-actin, which blocks further actin fiber polymerization. The current study showed that phosphorylated Gogo activates the actin-capping protein Hts to prevent uncontrolled actin polymerization in R8 axon termini. The overexpression of Hts in R8 axons alone did not prevent R8 filopodia extension, thus suggesting that phosphorylated Gogo is required. However, a previous cell culture study demonstrated that physical Gogo/Hts interactions take place regardless of the phosphorylation status of the YYD motif (Mann et al., 2012). This suggests that phosphorylated Gogo regulates Hts enzymatic activity rather than binding. The enzymatic activity of the Hts homolog adducin is controlled by Ser/Thr kinases in mammals (Fukata et al., 1999; Matsuoka et al., 1996; Matsuoka et al., 2000). This type of Ser/Thr kinase activation might occur in conjunction with the activation of the Tyr kinase that phosphorylates the Gogo YYD motif. These regulations may result in Gogo counteracting Fmi to suppress filopodia extension during step 2.

### Genomic economy of Gogo regulation in neuronal circuit formation

This study demonstrates that the insulin secreted from surface and cortex glia switches the phosphorylation status of Gogo, thereby regulating its two distinct functions. Nonphosphorylated Gogo mediates the initial recognition of the glial protrusion in the medulla column center. Phosphorylated Gogo suppresses horizontal filopodia extension by counteracting Fmi to maintain the one axon to one column ratio (Figure 7I).

Phosphorylated protein is typically activated or inactivated by phosphorylation. For example, to become activated and transduce downstream signaling, Robo and Eph have tyrosine phosphorylation sites and need to be dephosphorylated or phosphorylated, respectively (Dearborn et al., 2002; Sun et al., 2000). Few proteins have two distinct functions that are independently assigned to phosphorylation status (Li et al., 2018), and the current study demonstrates that Gogo is one of them. This mechanism is of great interest from a genomic economy point of view, where the animal genome takes an economical strategy to maximize protein functions and networks with a limited number of genes. The genomic economical strategy was likely important in the establishment of complex functional neuronal circuits during the evolution of higher-order species. Therefore, this mechanism is highly likely to be conserved across species.

## Acknowledgements

We gratefully acknowledge Dr. Pierre-Yves Plaçais (CNRS France) for providing the dlip4-Gal4 line and Dr. Thomas R Clandinin (Stanford Univ.) for FlpStop cassette. We thank Kyoto Stock Center (DGRC), Bloomington Drosophila Stock Center, Vienna Drosophila Resource Center (VDRC), and Developmental Studies Hybridoma Bank for providing fly or antibody stocks. We thank Enago (www.enago.jp) for the English language review. This work was supported by Grant-in-Aid for JSPS Fellows 19J14499 (H.T), 18J00367 (Y.N.), JSPS KAKENHI Grant number 18K06250 (S.H.-S.), 18K14835 (Y.N.), 17H03542 (M.S.), 17H04983 (A.S.), 19K22592 (A.S.), Grant-in Scientific Research on Innovation Areas from Ministry of Education, Culture, Sports, Science, and Technology of Japan “Dynamic regulation of Brain Function by Scrap & Build System” 16H06457 (T.S.), 17H05739 (M.S.) “Interplay of developmental clock and extracellular environment in brain formation” 17H05761 (M.S.), 19H04771 (M.S.), Takeda Science Foundation Life Science Research Grant (A.S) and Takeda Visionary Research Grant from the Takeda Science Foundation (T.S.).

## Author Contributions

Conceptualization, H.T., S.H-S., Y.N., A.S., and T.S.; Methodology, H.T., S.H-S., Y.N., A.S., M.S., and T.S.; Investigation, H.T., S.H-S., Y.N., Y.I.; Writing – Original Draft, H.T., S.H-S., Y.N., and T.S.; Writing – Review & Editing, H.T., S.H-S., Y.N., A.S. and T.S.; Funding Acquisition, H.T., S.H-S., Y.N., A.S., M.S., and T.S.; Supervision, A.S., M.S., and T.S.

