## Supplementary Data for "Glial insulin regulates cooperative or antagonistic Golden goal/Flamingo interactions during photoreceptor axon navigation"

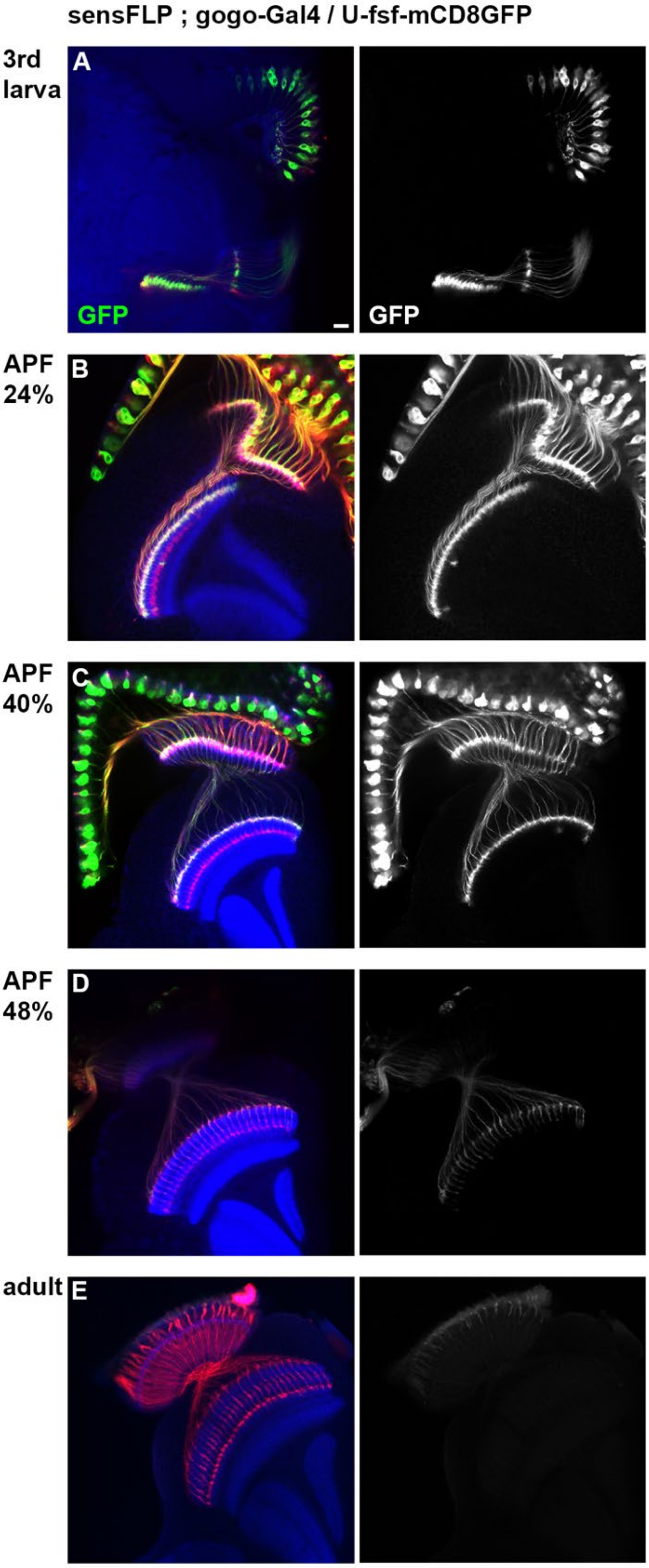

**Supplementary figure 1.** ***gogo*** **expression gradually declines during midpupal stages**

(A-E) *gogo* expression level in R8 cells are monitored by *gogo*-Gal4 sensFLP UAS-FsF-mCD8GFP. *gogo*-Gal4 is knocked into the *gogo* intron locus by MiMIC system. Photoreceptor axons are labeled with 24B10 (red) and medulla layers with anti-N-cadherin (blue). The GFP protein was strongly observed at 3$^{rd}$ larval stage (A) and APF 24% (B), then gradually declined during midpupal stages (APF40% (C) and 48% (D)). Scale bar :10μm.

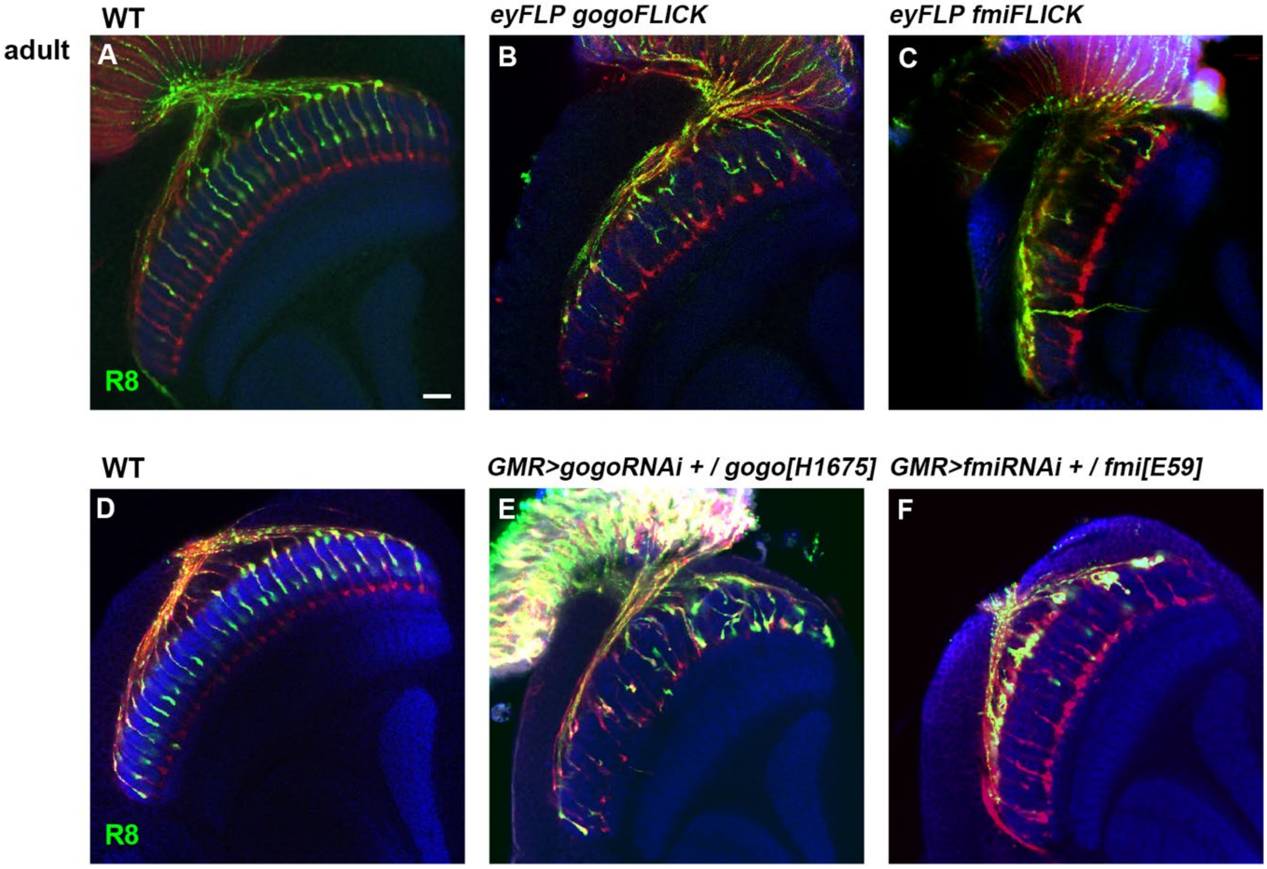

**Supplementary figure 2. R cells specific null mutant of Gogo and Fmi**

(A-F) R cells of control (A and D) and *gogo* (B and E), *fmi* (C and F) mutants in adult visualized with GFP (green) counterstained with 24B10 (red) and N-Cadherin (blue). *gogo* and *fmi* heterozygote mutant with R cell specific RNAi (GMR-Gal4, UAS-RNAi, at 29℃) (E and F respectively) showed strong phenotype equivalent to *gogo, fmi eyFLICK* flies (*gogo*[H1675]/<*gogo*<, *fmi*[E59]/<*fmi*<[2]) (B and C respectively) in adult. Scale bar:10μm.

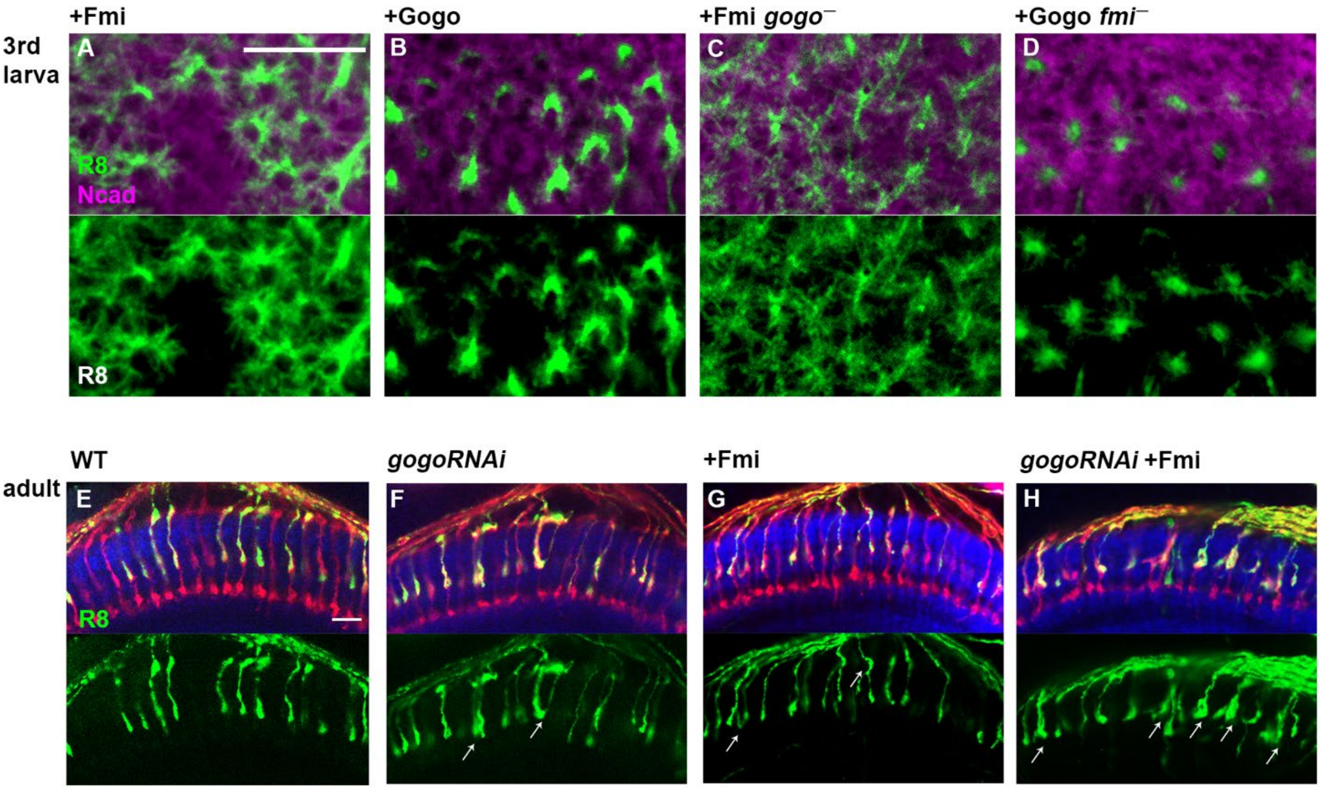

**Supplementary figure 3. Gogo and Fmi functions are not redundant**

(A-D) Mutual rescue between *gogo* and *fmi* was tested. R8 specific *gogo* and *fmi* mutant clones were generated by heterozygote mutation with R8 specific RNAi. R8 specific expression of Gogo or Fmi was driven by the sensFLP, GMR-FsF-Gal4. R8 axons are visualized with mCD8GFP (green), and counterstained with N-Cadherin (magenta). Overexpression of Gogo or Fmi in the other mutant background did not show any mutual rescue (C-D).

(E-H) The expression of Gogo was downregulated using RNAi or upregulated by overexpression of transgenes in Fmi. The RNAi and transgenes were expressed under the sensFLP GMR-FsF-Gal4 driver. R8 axons are labeled with UAS-mCD8GFP (green), and counterstained with mAb24B10 (red) and N-Cadherin (blue). In wild type, R8 axons do not bundle each other and target M3 layer. In Fmi overexpression or gogo knock-down, the R8 axons bundling phenotype (arrows in F-G) is shown in the adult stage, and it was also promoted by gogo RNAi (arrows in H). Scale bars:10μm.

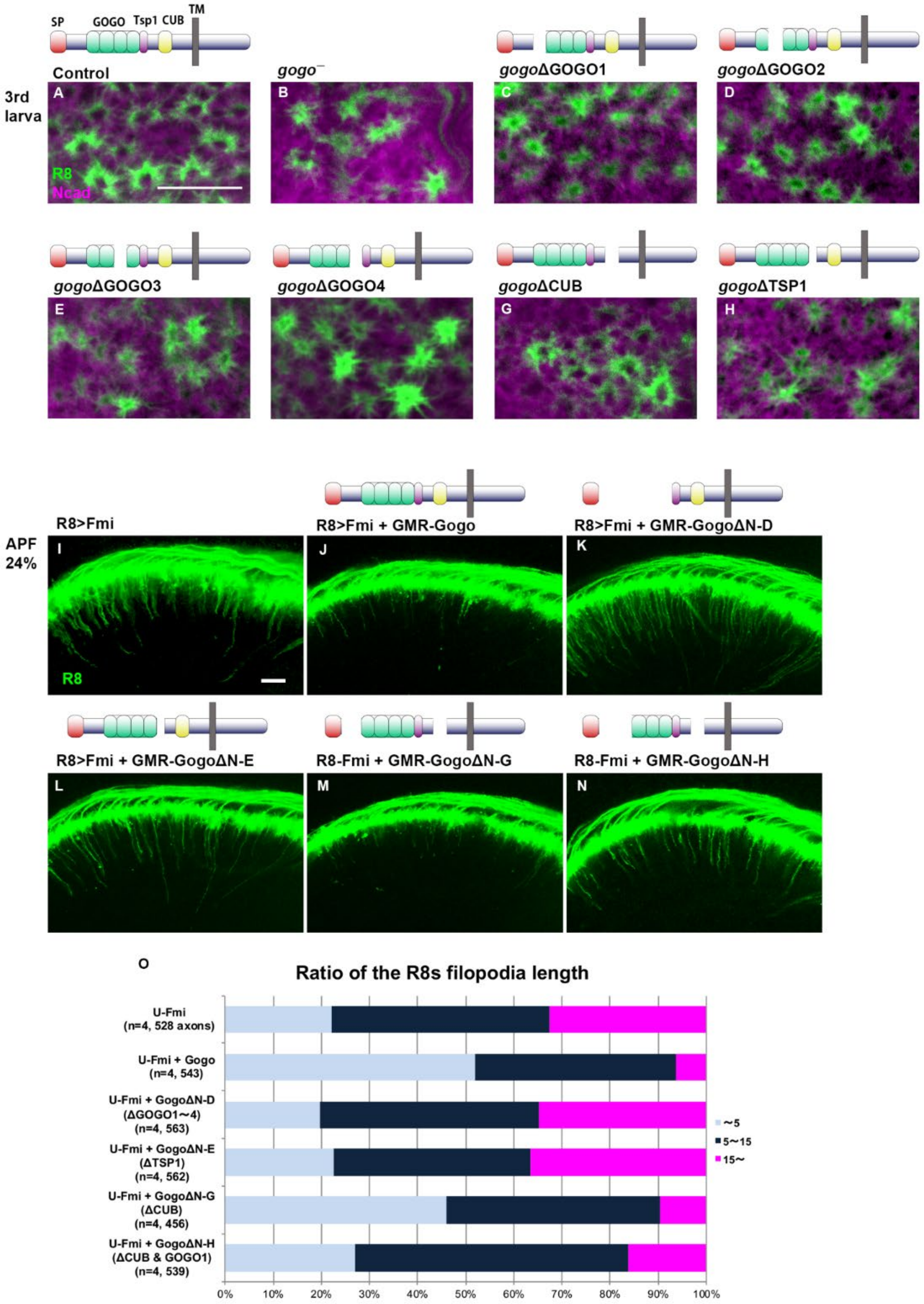

**Supplementary figure 4. Gogo ecto-domain interacts with Fmi**

(A-H). R8 axons are labeled with myr-Tomato (green) counterstained with N-Cadherin (magenta). The illustrations above the images show the structures of the Gogo extracellular deletion series. Small deletions of GOGO (C-F) or Tsp1 (H) domains heterozygous with *gogo* null mutation resulted in the R8 axons targeting defects equivalent to gogo null mutant (B) at 3rd larval stage.

(I-O) R8 axons are labeled with mCD8GFP (green) in Fmi overexpression. The transgenes were expressed in all photoreceptor neurons by GMR promoter. Gogo which lacks GOGO domain (k and N) or Tsp1 domain (L) showed weaker suppression of filopodia extension phenotype in Fmi overexpression than wild type Gogo (J). (O) Quantification of R8 axons filopodia length. The longest filopodia in a 3D image was measured from one axon. The length of filopodia is divided into 3 classes of ~ 5μm (Light blue), 5-15μm (dark blue), 15μm ~(magenta). Scale bars: 10μm.

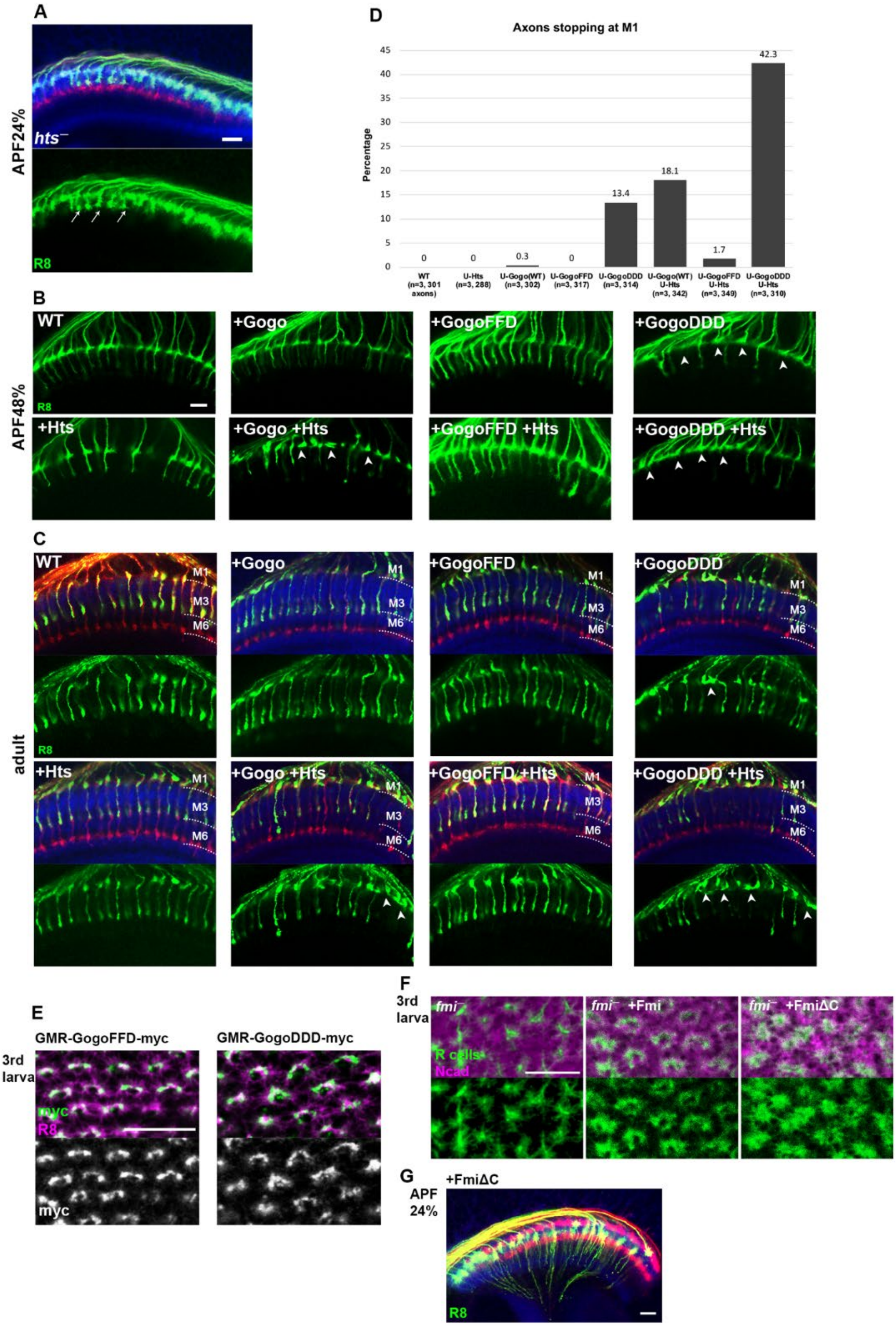

**Supplementary figure 5. Gogo and Fmi cytoplasmic domain change its functional properties**

(A) R8 specific *hts* mutant were generated by *hts* heterozygote with R8 specific RNAi. In APF 24% and adult stage, *hts* mutant show R8 axons bundling phenotype (arrows) due to the excessive extension of filopodia in random direction at APF 24% stage.

(B-D) Phenotypes of R8 axons overexpressing Gogo (wild type, phosphorylated, non-phosphorylated) and Hts. Transgenes were expressed under the GMR-Gal4 driver (25°C). R8 axons are labeled with sensFLP UAS-FsF-mCD8GFP (green), and counterstained with mAb24B10 (red) and N-Cadherin (blue). When Gogo, non-phospho-Gogo or Hts were overexpressed alone, the R8 axons target normally. In phospho-Gogo overexpression, the filopodia extension was partially suppressed at APF 48% (B, arrow) and stop at the medulla neuropil surface in adult (C, arrow).

Overexpression of Hts in combination with Gogo or phospho-Gogo, but not with non-phospho-Gogo enhanced the suppression of the filopodia extension and R8 axons stopped at the M1 layer (B-C arrowheads). (D) Quantification of R8 axons stopping at M1 layer in adult.

(E) The myc tagged GMR *gogo* transgenes were expressed in all photoreceptor neurons and detected by anti-myc (green). R8 cells labeled with *sens*-Gal4, UAS-mCD8GFP (magenta). The localization of Gogo dephospho-mimetic and phospho-mimetic versions did not differ in R8 axon termini at the 3rd larval stage.

(F) The requirement of the fmi cytoplasmic part was confirmed in the rescue experiments. *fmi* mutant clones were generated by ey3.5FLP; <*fmi*< [2] / *fmi*[E59]. Expression of UAS-Fmi constructs was driven by the GMR-Gal4 driver. R axons are visualized with ey3.5FLP, UAS-FsF-mCD8GFP (green), and counterstained with N-Cadherin (magenta). Wild-type Fmi but not FmiΔC rescued the *fmi* mutant phenotype in 3rd larval stage.

(G) FmiΔC was expressed under the sensFLP GMR-FsF-Gal4. R8 axons are labeled with UAS-mCD8GFP (green), and counterstained with mAb24B10 (red) and N-Cadherin (blue). In FmiΔC overexpression, R8 cells extend their vertical filopodia precociously towards the deeper layer of the medulla at APF 24%, similar to wild-type Fmi overexpression. Scale bars :10μm.

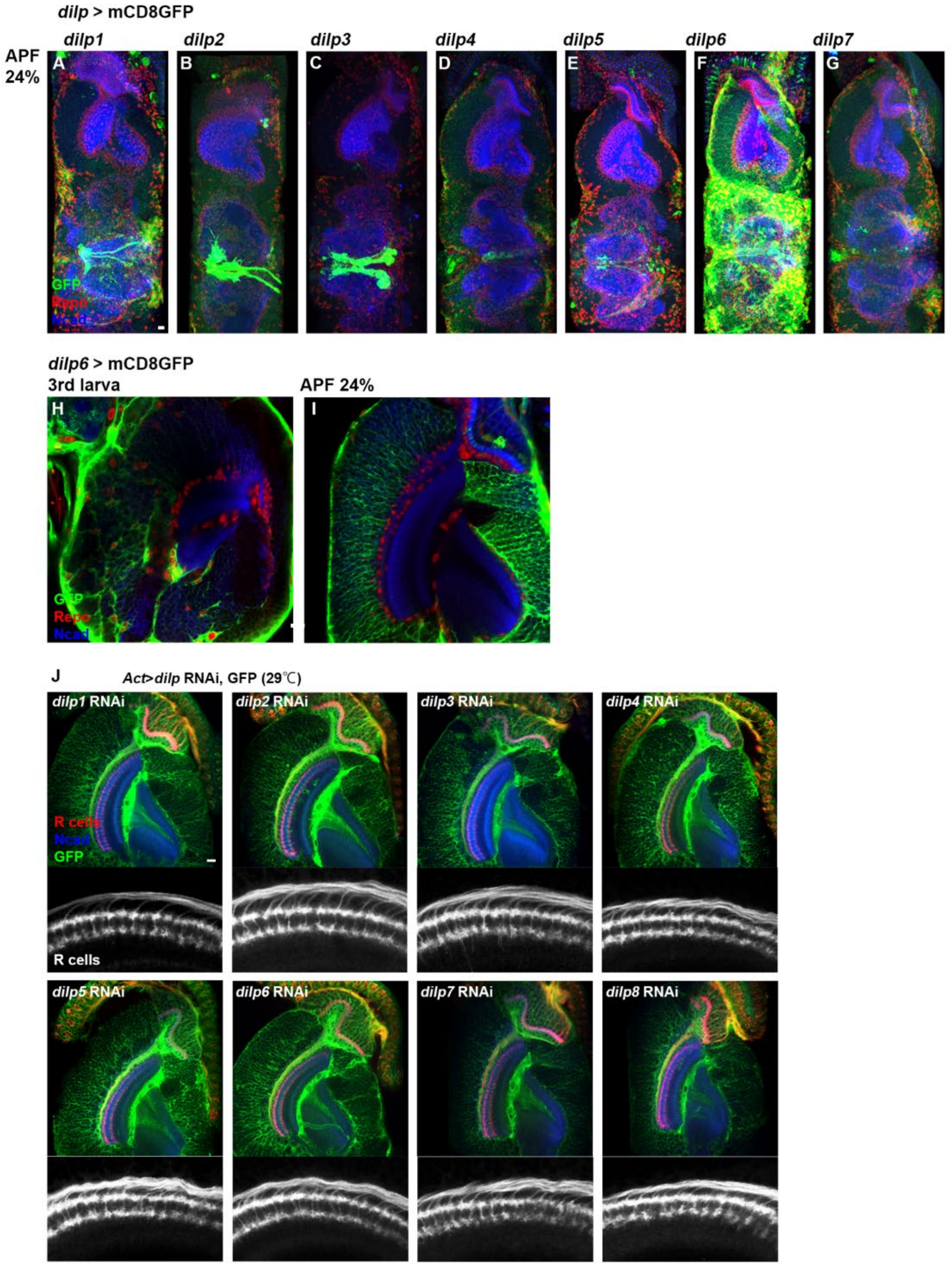

**Supplementary figure 6. *dilp* genes expression pattern in optic lobe**

(A-I) *dilp* gene expression was monitored by *dilp*-Gal4, UAS-mCD8GFP (green) and counterstained with Repo (red) and N-Cadherin (blue). *dilp*1, 2, 3, 5 are strongly express in insulin producing cells (IPCs) (A-C and E). *dilp6, dilp4* and *dilp7* were broadly expressed in glial cells (D, F and G), especially *dilp6* strongly express in cortex and surface glia in the optic lobe (F, H and I). While entire structure of the brain changes from 3rd instar larvae to APF 24%, position of the glia cells also changes. Cortex glia does not cover medulla neuropil in $3^{rd}$ larval stage (H), but does in APF24% (I). (J) *dilp* genes were knocked down in the whole animal using Act-Gal4, UAS-*dilp* RNAi, tub-Gal80$^{ts}$. RNAi positive cells are labeled with UAS-mCD8GFP (green), and counterstained with mAb24B10 (red) and N-Cadherin (blue). In all knockdowns of dilp1-8, R8 axons did not bundle each other and showed any phenotype compared to the WT. Scale bars :10μm.

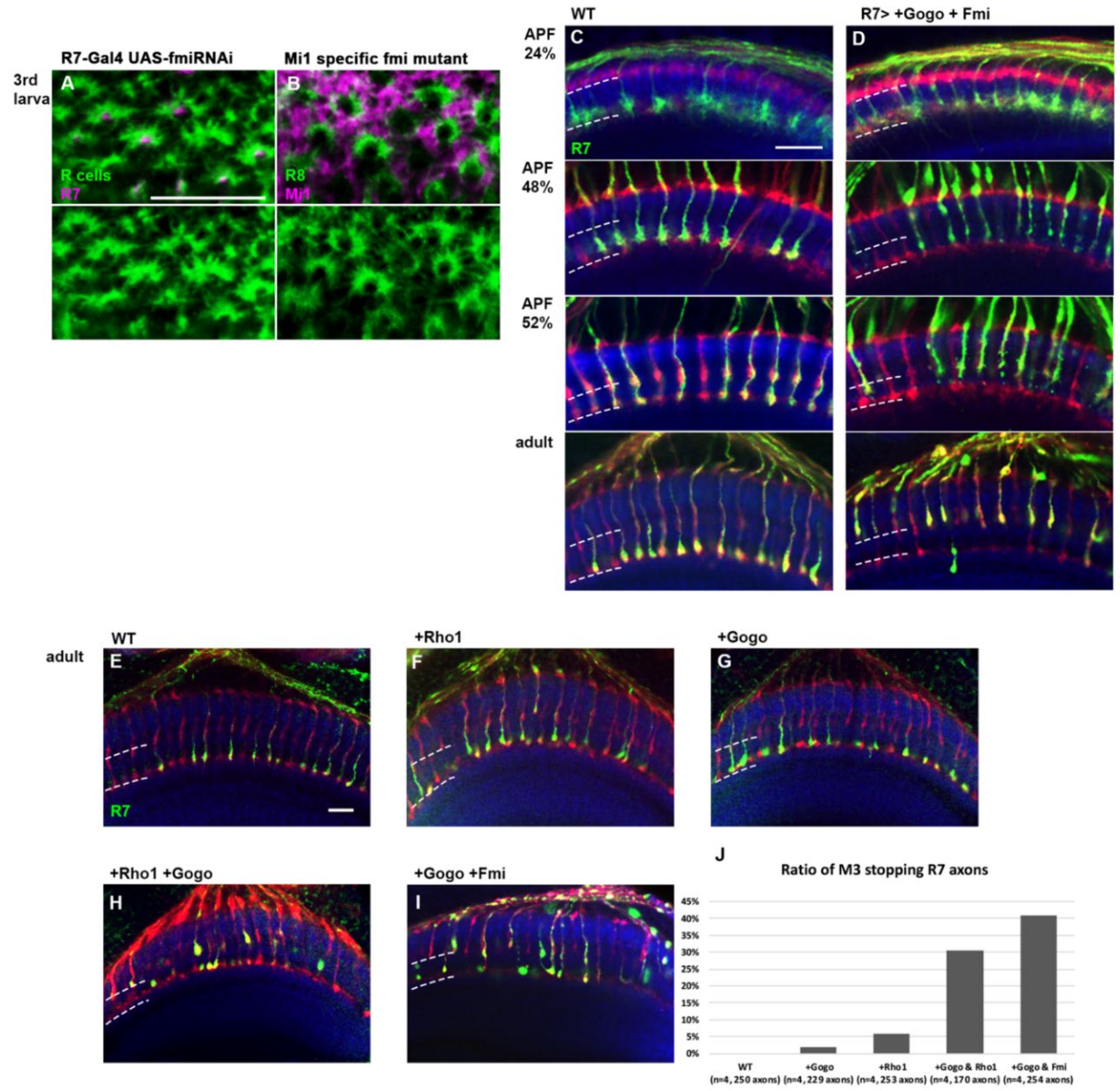

**Supplementary figure 7. Gogo and Fmi interact with Fmi to regulate cytoskeletal reorganization**

(A) In 3rd larval stage, R7 neurons that is known to be the first core member of the medulla column formation were labeled with 20C11FLP, GMR-FsF-Gal4, UAS-mCD8GFP (magenta), and R axons with 24B10 (green). R7 specific *fmi* knockdown did not show any notable defect in overall R8 axon targeting or morphology of the R8 axon termini.

(B) Mi1 neurons labeled with bshM-Gal4, UAS-mCD8GFP (magenta) and R8 axons with myrTomato (green). In Mi1 specific *fmi* knockdown, no differences were seen in R8 axons from that of WT.

(C–D) Gogo and Fmi were co-overexpressed in R axons using GMR-Gal4 during pupal stage. R7 axons were visualized with mCD8GFP (green) and counterstained with mAb24B10 (red) and N-Cadherin (blue). In APF 24%, R7 axons with co-overexpression reached the R7 temporary layer correctly. From APF 48% to 52%, R7 growth cones collapsed and retraction of R7 axons was observed. Almost all the R7 axons retracted and stopped at M3 layer in adult.

(E-J) The R7 axons of control (E) and overexpression using GMR-Gal4, UAS-*gogo* and/or GMR-Rho1 were visualized with Rh4-GFP (green) and counterstained with mAb24B10 (red) and N-Cadherin (blue). R7 photoreceptor axons targeted normally when Rho1 (F) or Gogo (G) were overexpressed alone. Co-overexpression of Gogo and Rho1 caused R7 photoreceptor mistargeting to the M3 layer (H), similar to co-overexpression of Gogo and Fmi (I). The quantification of the R7 photoreceptor axon mistargeting to M3 layer is shown in (J). Scale bars :10μm.

**Supplementary Table 1 List of all full genotypes used**

| Figure | Genotype (Genotypes unrelated to the context are in brackets.) |
|---|---|
| 1C | sensFLP / UAS-FLP; GMR-FsF-Gal4, UAS-myrRFP/+; gogo-FsF-GFP, sensGal4 / gogo-FsF-GFP |
| 1D-1G | sensFLP ; gogo-FsF-GFP |
| 1H | sensFLP ; fmi-FsF-mcherry ; sensGal4, UAS-mCD8GFP / + |
| 1I-1L | sensFLP ; fmi-FsF-mcherry |
| 1M-1N | sensFLP ; fmi-FsF-mcherry ; gogo-FsF-GFP |
| 2A-2C | sensFLP ; GMR-FsF-Gal4/+; UAS-mCD8GFP / + |
| 2D-2F | sensFLP ; GMR-FsF-Gal4/UAS-gogoRNAi (GD3616) ; UAS-mCD8GFP / gogo[H1675] |
| 2H-2J | sensFLP ; GMR-FsF-Gal4/fmi[E59]; UAS-mCD8GFP / UAS-fmiRNAi (VDRC; GD607) |
| 3A | sensFLP ; GMR-FsF-Gal4/UAS-gogoRNAi (GD3616) ; UAS-mCD8GFP / + |
| 3B | sensFLP ; GMR-FsF-Gal4/+ ; UAS-mCD8GFP / UAS-fmiRNAi (VDRC; GD607) |
| 3C | sensFLP ; GMR-FsF-Gal4/gogoRNAi (GD3616) ; UAS-mCD8GFP / UAS-fmiRNAi (VDRC; GD607) |
| 3E | sensFLP ; GMR-FsF-Gal4/+; UAS-mCD8GFP / UAS-Gogo (V) |
| 3F | sensFLP ; GMR-FsF-Gal4/+; UAS-mCD8GFP / UAS-Fmi |
| 3G | sensFLP ; GMR-FsF-Gal4/UAS-gogoRNAi (GD3616); UAS-mCD8GFP / UAS-Fmi |
| 3H | sensFLP ; GMR-FsF-Gal4/UAS-Gogo (V); UAS-mCD8GFP / UAS-Fmi |
| 4A | Same as 1C |
| 4B | sensFLP / UAS-FLP; GMR-FsF-Gal4, UAS-myrRFP/fmi[E59]; gogo-FsF-GFP, sensGal4 / gogo-FsF-GFP, UAS-fmiRNAi (VDRC; GD607) |
| 4C | sensFLP / UAS-FLP; GMR-FsF-Gal4, UAS-myrRFP/+; gogo-FsF-GFP, sensGal4 / gogo-FsF-GFP, UAS-Fmi |
| 4D | Same as 1C |
| 4E | Same as 4C |
| 4F | Same as 1H |
| 4G | sensFLP ; fmi-FsF-ncherry / fmi-FsF-mcherry, UAS-gogoRNAi(GD3616) ; sensGal4, UAS-mCD8GFP / gogo[H1675] |
| 4H | sensFLP ; fmi-FsF-mcherry ; sensGal4, UAS-mCD8GFP / UAS-Gogo (V) |
| 4I | Same as 1C |
| 4J | Same as 4C |
| 5A | sensFLP ; GMR-Gal4/UAS-FsF-mCD8GFP; gogo[H1675] / gogo[D1600] |
| 5B | sensFLP ; GMR-Gal4, UAS-GogoFL/UAS-FsF-mCD8GFP; gogo[H1675] / gogo[D1600] |

| | |
|---|---|
| 5C | sensFLP ; GMR-Gal4, UAS-FsF-mCD8GFP/ UAS-GogoΔC; gogo[H1675] / gogo[D1600] |
| 5D | sensFLP ; GMR-Gal4, UAS-GogoFFD/UAS-FsF-mCD8GFP; gogo[H1675] / gogo[D1600] |
| 5E | sensFLP ; GMR-Gal4, UAS-GogoDDD/UAS-FsF-mCD8GFP; gogo[H1675] / gogo[D1600] |
| 5G | sensFLP; GMR-Gal4 / + ; UAS-FsF-mCD8GFP / UAS-Fmi |
| 5H | sensFLP; GMR-Gal4 / UAS-GogoFL; UAS-FsF-mCD8GFP / UAS-Fmi |
| 5I | sensFLP; GMR-Gal4 / UAS-GogoΔC; UAS-FsF-mCD8GFP / UAS-Fmi |
| 5J | sensFLP; GMR-Gal4 / UAS-GogoFFD; UAS-FsF-mCD8GFP / UAS-Fmi |
| 5K | sensFLP; GMR-Gal4 / UAS-GogoDDD; UAS-FsF-mCD8GFP / UAS-Fmi |
| 5M | Same as 5A |
| 5N | Same as 5B |
| 5O | Same as 5D |
| 6A | sensFLP; GMR-FsF-Gal4/dinr[ex15]; UAS-mCD8GFP / dinrRNAi (BDSC 31037) |
| 6B | y dilp6-Gal4/+; UAS-nlsGFP/+ |
| 6C, 6D | yw; senslexA lexAopTomato/ato-t-myc ; loco-Gal4, UAS-mCD8GFP /UAS-shi[ts1] |
| 6E | yw; senslexA lexAopTomato/ato-t-myc ; R85G01-Gal4, UAS-mCD8GFP /UAS-shi[ts1] |
| 6F | yw; senslexA lexAopTomato/ato-t-myc ; R25A01-Gal4, UAS-mCD8GFP /UAS-shi[ts1] |
| 6G | yw; senslexA lexAopTomato/Mz97-Gal4, UAS-stinger; ato-t-myc /UAS-shi[ts1] |
| 6H | y dilp6-Gal4; senslexA lexAopTomato/+ ; + /UAS-shi[ts1] |
| 6I | yw; senslexA lexAopTomato/dilp2-Gal4 ; + /UAS-shi[ts1] |
| 6K | yw; senslexA lexAopTomato/hobRNAi (BDSC 66966); loco-Gal4, UAS-mCD8GFP /+ |
| 7A, 7C | yw; senslexA lexAopTomato/+ ; loco-Gal4, UAS-mCD8GFP /+ |
| 7B | UAS-FLP/yw; fmi-FsF-mcherry; loco-Gal4, UAS-mCD8GFP/+ |
| 7D | yw; senslexA lexAopTomato/fmi[E59]; loco-Gal4, UAS-mCD8GFP / UAS-fmiRNAi (VDRC; GD607) |
| 7E | sensFLP; fmi-FsF-mcherry/+; locoGal4, UAS-mCD8GFP/+ |
| 7F | sensFLP; fmi-FsF-mcherry / fmi[E59] ; locoGal4, UAS-mCD8GFP/ UAS-fmiRNAi (VDRC; GD607) |
| 7G | sensFLP; ; locoGal4/ gogo-FsF-GFP |
| 7H | sensFLP; fmi[E59] / +; locoGal4/ gogo-FsF-GFP, UAS-fmiRNAi (VDRC; GD607) |
| 8B | w;UAS-mCD8GFP/+;; OK107-Gal4/+ |
| 8C | w; UAS-mCD8GFP/+; gogo[H1675] / gogo[D1600]; OK107-Gal4/+ |
| 8D | w; UAS-mCD8GFP/UAS-gogoFL; gogo[H1675] / gogo[D1600]; OK107-Gal4/+ |
| 8F | w, UAS-dicer2/+;UAS-mCD8GFP/UAS-fmiRNAi;;OK107-Gal4/+ |
| 8G | w, UAS-dicer2/+;UAS-mCD8GFP/+;UAS-Fmi/+;OK107-Gal4/+ |
| 8H | w;UAS-dicer/UAS-fmiRNAi;repo-Gal4/+; |

| | |
|---|---|
| control in 8G | w, UAS-dicer2/+;UAS-mCD8GFP/40D-UAS;;OK107-Gal4/+ |
| gogo RNAi in 8H | w, UAS-dicer2/+;UAS-mCD8GFP/UAS-gogoRNAi;;OK107-Gal4/+ |
| gogo RNAi Fmi OE in 8H | UAS-dicer2/+;UAS-mCD8GFP/UAS-gogoRNAi;UAS-Fmi/+;OK107-Gal4/+ |
| S1 | sensFLP ; gogo-Gal4 / UAS-FsF-mCD8GFP |
| S2A | Rh6-GFP, eyFLP |
| S2B | Rh6-GFP, eyFLP; <gogo< /gogo[H1675] |
| S2C | Rh6-GFP, eyFLP; <fmi< / fmi[E59] |
| S2D | sensFLP ; GMR-Gal4/ + ; UAS-FsF-mCD8GFP / + |
| S2E | sensFLP ; GMR-Gal4/UAS-gogoRNAi(GD3616) ; UAS-FsF-mCD8GFP / gogo[H1675] |
| S2F | sensFLP; GMR-Gal4/fmi[E59] ; UAS-FsF-mCD8GFP / UAS-fmiRNAi |
| S3A | Same as 3F |
| S3B | Same as 3E |
| S3C | sensFLP ; GMR-FsF-Gal4, UAS-FsF-mCD8GFP / UAS-gogoRNAi(GD3616) ; UAS-Fmi / gogo[H1675] |
| S3D | sensFLP ; GMR-FsF-Gal4, UAS-FsF-mCD8GFP / fmi[E59] ; UAS-Gogo (V) / UAS-fmiRNAi (VDRC; GD607) |
| S3A | Same as 2A |
| S3B | Same as 3A |
| S3C | Same as 3F |
| S3D | Same as 3G |
| S4A | yw; senslexA lexAopTomato/+ ; gogo[H1675] /+ |
| S4B | yw; senslexA lexAopTomato/+ ; gogo[H1675] /gogo[D1600] |
| S4C | yw; senslexA lexAopTomato/+ ; gogo[H1675] /gogoΔGOGO1 |
| S4D | yw; senslexA lexAopTomato/+ ; gogo[H1675] /gogoΔGOGO2 |
| S4E | yw; senslexA lexAopTomato/+ ; gogo[H1675] /gogoΔGOGO3 |
| S4F | yw; senslexA lexAopTomato/+ ; gogo[H1675] /gogoΔGOGO4 |
| S4G | yw; senslexA lexAopTomato/+ ; gogo[H1675] /gogoΔCUB |
| S4H | yw; senslexA lexAopTomato/+ ; gogo[H1675] /gogoΔTSP1 |
| S4I | Same as 3F |
| S4J | sensFLP ; GMR-FsF-Gal4/GMR-GogoFL; UAS-mCD8GFP / UAS-Fmi |
| S4K | sensFLP ; GMR-FsF-Gal4/GMR-GogoΔN-D; UAS-mCD8GFP / UAS-Fmi |
| S4L | sensFLP ; GMR-FsF-Gal4/GMR-GogoΔN-E; UAS-mCD8GFP / UAS-Fmi |
| S4M | sensFLP ; GMR-FsF-Gal4/GMR-GogoΔN-G; UAS-mCD8GFP / UAS-Fmi |
| S4N | sensFLP ; GMR-FsF-Gal4/GMR-GogoΔN-H; UAS-mCD8GFP / UAS-Fmi |
| S5 | sensFLP ; GMR-FsF-Gal4/FRT42D hts null; UAS-mCD8GFP / htsRNAi (BDSC 35421) |
| S5B, S5C | sensFLP; GMR-Gal4 / + ; UAS-FsF-mCD8GFP / + |
| | sensFLP; GMR-Gal4, UAS-GogoFL / + ; UAS-FsF-mCD8GFP / + |

| | |
|---|---|
| | sensFLP; GMR-Gal4, UAS-GogoFFD / + ; UAS-FsF-mCD8GFP / + |
| | sensFLP; GMR-Gal4, UAS-GogoDDD / + ; UAS-FsF-mCD8GFP / + |
| | sensFLP; GMR-Gal4 / + ; UAS-FsF-mCD8GFP / UAS-Add1-myc |
| | sensFLP; GMR-Gal4 / UAS-GogoFL ; UAS-FsF-mCD8GFP / UAS-Add1-myc |
| | sensFLP; GMR-Gal4 / UAS-GogoFFD ; UAS-FsF-mCD8GFP / UAS-Add1-myc |
| | sensFLP; GMR-Gal4 / UAS-GogoDDD ; UAS-FsF-mCD8GFP / UAS-Add1-myc |
| S5E | GMR-gogoFFD T4a / + ; sens-Gal4, UAS-mCD8GFP / + |
| | GMR-gogoDDD T1a / + ; sens-Gal4, UAS-mCD8GFP / + |
| S5F | ey3.5FLP ; <fmiN< / fmi[E59], GMR-Gal4; UAS-FsF-mCD8GFP / + |
| | ey3.5FLP ; <fmiN< / fmi[E59], GMR-Gal4; UAS-FsF-mCD8GFP / UAS-Fmi |
| | ey3.5FLP ; <fmiN< / fmi[E59], GMR-Gal4; UAS-FsF-mCD8GFP / UAS-FmiΔC |
| S5H | sensFLP ; GMR-FsF-Gal4 / + ; UAS-mCD8GFP / UAS-FmiΔC |
| S6A | dilp1-Gal4 / + ; UAS-mCD8GFP / + |
| S6B | dilp2-Gal4 / + ; UAS-mCD8GFP / + |
| S6C | dilp3-Gal4 / + ; UAS-mCD8GFP / + |
| S6D | dilp4-Gal4 / + ; UAS-mCD8GFP / + |
| S6E | dilp5-Gal4 / + ; UAS-mCD8GFP / + |
| S6F, S6H, S6I | y, dilp6-Gal4 / + ; UAS-mCD8GFP / + |
| S6G | UAS-mCD8GFP / dilp7-Gal4 |
| S6J | Act-Gal4, UAS-mCD8GF / + ; tub-Gal80ts/ dilp1RNAi |
| | Act-Gal4, UAS-mCD8GF / + ; tub-Gal80ts/ dilp2RNAi |
| | Act-Gal4, UAS-mCD8GF / + ; tub-Gal80ts/ dilp3RNAi |
| | Act-Gal4, UAS-mCD8GF / + ; tub-Gal80ts/ dilp4RNAi |
| | Act-Gal4, UAS-mCD8GF / + ; tub-Gal80ts/ dilp5RNAi |
| | Act-Gal4, UAS-mCD8GF / + ; tub-Gal80ts/ dilp6RNAi |
| | Act-Gal4, UAS-mCD8GF / + ; tub-Gal80ts/ dilp7RNAi |
| | Act-Gal4, UAS-mCD8GF / dilp8RNAi ; tub-Gal80ts/ + |
| S7A | 20C11FLP/GMR-FsF-Gal4 ; UAS-mCD8GFP / UAS-fmiRNAi (VDRC; GD607) |
| S7B | senslexA lexAopTomato/fmi[E59] ; bshM-Gal4, UAS-myrGFP (from Kanazawa uni) / UAS-fmiRNAi (VDRC; GD607) |
| S7C | 20C11FLP/GMR-FsF-Gal4 ; UAS-mCD8GFP / + |
| S7D | 20C11FLP/GMR-FsF-Gal4, U-gogo (V) ; UAS-mCD8GFP / U-Fmi |
| S7E | Rh4-GFP/+ |
| S7F | Rh4-GFP/+ ; GMR-Gal4, U-Gogo(V)/+ |
| S7G | Rh4-GFP/+ ; GMR-RhoA/+ |
| S7H | Rh4-GFP/+ ; GMR-Gal4, U-Gogo(V)/+ ; GMR-RhoA/+ |
| S7I | 20C11FLP/GMR-Gal4, U-gogo (V) ; UAS-FsF-mCD8GFP / U-Fmi |
| Movie gogo mutant | sensFLP ; GMR-Gal4 / UAS-FsF-mCD8GFP ; gogo-Flpstop / gogo[H1675] |
| Movie control | sensFLP ; GMR-Gal4 / UAS-FsF-mCD8GFP ; + / gogo[H1675] |

**Supplementary Table 2 oligo DNAs used for generating and analyzing transgenic flies**

| | |
|---|---|
| CTTCGCAGAATATACCGCTCTTCC | Gogo gDNA1 GogoΔGOGO1 |
| AAACGGAAGAGCGGTATATTCTGC | Gogo gDNA1 GogoΔGOGO1 |
| CTTCGGAGCTAATTTACTGCGGCA | Gogo gDNA2 GogoΔGOGO1, ΔGOGO2 |
| AAACTGCCGCAGTAAATTAGCTCC | Gogo gDNA2 GogoΔGOGO1, ΔGOGO2 |
| CTTCGATTCCCGTGTGCGGATCAC | Gogo gDNA3 GogoΔGOGO2, ΔGOGO3 |
| AAACGTGATCCGCACACGGGAATC | Gogo gDNA3 GogoΔGOGO2, ΔGOGO3 |
| CTTCGCAGTACTCTACGTACTTCT | Gogo gDNA4 GogoΔGOGO3, ΔGOGO4 |
| AAACAGAAGTACGTAGAGTACTGC | Gogo gDNA4 GogoΔGOGO3, ΔGOGO4 |
| CTTCGCCACCGGTATCCGAGTCTG | Gogo gDNA5 GogoΔTSP1 |
| AAACCAGACTCGGATACCGGTGGC | Gogo gDNA5 GogoΔTSP1 |
| CTTCGTAAATGCAGTCCCACGTGT | Gogo gDNA6 GogoΔTSP1, ΔCUB |
| AAACACACGTGGGACTGCATTTAC | Gogo gDNA6 GogoΔTSP1, ΔCUB |
| CTTCGTTCTCTGCCGGCGATGGAC | Gogo gDNA7 GogoΔCUB |
| AAACGTCCATCGCCGGCAGAGAAC | Gogo gDNA7 GogoΔCUB |
| CTTCGCAGTACTCTACGTACTTCT | Gogo gDNA8 GogoΔGOGO4 |
| AAACAGAAGTACGTAGAGTACTGC | Gogo gDNA8 GogoΔGOGO4 |
| AAGCTGAAAAGTGCGAGAATGTCAAG | FW gogoFlpstop attB-UAS Tom |
| CATGGACTTGGACATCTGAGTGTTTG | REV gogoFlpstop attB-UAS Tom |
| TATAGCGGCCGCATTGAAGTTCCTATTCCGAAGTTCC | FW FRT |
| TATAACTAGTCAAAAGCGCTCTGAAGTTCCTATAC | REV FRT |
| TATAACTAGTTAAATCCAGACATGATAAGATACATTGATGAG | FW stop-FRT |
| TATACTGCAGCAAAAGCGCTCTGAAGTTCCTATAC | REV stop-FRT |
| GCGCCTCGAGAACTTCGTATAGCATACATTA | FW loxP-RFP-loxP |
| TATACTCGAGAACGTGTCGGTACCAATTGAGCTC | REV loxP-RFP-loxP |
| TATAGCATGCTGCACTGGACATCATTGAACTT | FW mini-white |
| TATAGAATTCCCAGTGAAATCCAAGCATTTTCTA | REV mini-white |
| TATAAAGCTTGGATCCGGCTTACCTTATCTGG | FW GogoFsFGFP pre |
| TATAGCTAGCCACGGCGACTTCCTTTGACTTC | REV GogoFsFGFP pre |
| TATACTCGAGCACCAAGATATAATTGTACATAAAACC | FW GogoFsFGFP post |
| TATAGGTACCTACAGGTCGGGGTGATATAGAAA | REV GogoFsFGFP post |
| TATACTGCAGATGAGTAAAGGAGAAGAACTTTTC | FW GFP |
| TATACTCGAGTCTAGTGGATCCAGACATGATAAG | REV GFP |
| CTTCGGAGCCGAAGTCAAAGGAAG | gogo gDNA Gogo-FsF-GFP |
| AAACTTCCTTTGACTTCGGCTCC | gogo gDNA Gogo-FsF-GFP |
| TATAAAGCTTAGCAGCACCACAACAAAATCAA | FW fmiFsFmcherry pre |
| TATAGCTAGCATATTCCGCCTCTGAGTCGGTAT | REV fmiFsFmcherry pre |
| TATACTCGAGAAAAGGTCTGCAGCAAGATTGTCC | FW fmiFsFmcherry post |
| TATAGGTACCCCATAGCATTTTGCATTACGTCGAA | REV fmiFsFmcherry post |
| TATAGTCGACATGGTGAGCAAGGGCG | FW mcherry |

| | |
|---|---|
| TATACTCGAGTCTAGTGGATCCAGACATGATAAG | REV mcherry |
| CTTCGACCTTTTGGCCAACTTACT | Fmi gDNA Fmi-FsF-mCherry |
| AAACAGTAAGTTGGCCAAAAGGTC | Fmi gDNA Fmi-FsF-mCherry |
